# Collective fate decisions and cell rearrangements underlie gastruloid symmetry breaking

**DOI:** 10.1101/2024.12.16.628776

**Authors:** David Oriola, Gabriel Torregrosa-Cortés, Sotiris Samatas, Krisztina Arató, David Fernández-Munuera, Elisa Maria Hahn, Kerim Anlaş, Jordi Garcia-Ojalvo, Vikas Trivedi

## Abstract

How cell fate decisions coordinate with tissue-scale morphogenesis remains a major challenge in developmental biology. Gastruloids, three-dimensional aggregates of pluripo-tent stem cells that self-organise and break symmetry via polarised Brachyury/T expres-sion, provide an ideal system to address this question. By generating gastruloids with defined initial proportions of T-expressing cells we show that fate decisions occur collec-tively, with cell-fate proportions influencing the transition rates. Mechanical measure-ments reveal differences in surface tension between T-positive and T-negative tissues, con-sistent with radial cell sorting. Finally, incorporating fate dynamics and mechanics into a computational model recapitulates the sequential symmetry-breaking events observed in vitro. Our findings identify a mechanochemical mechanism underlying axis formation, and demonstrate how multicellular systems can robustly self-organise without external signalling cues.

## Introduction

How animals establish the body plan is a fundamental problem in biology (*1, 2*). Molecular cues from extra-embryonic tissues polarise the early embryo thus generating a coordinate sys-tem for subsequent regional patterning (*3–7*). In the last decade, several studies have shown that collections of stem cells can break symmetry *in vitro* even in the absence of instructive signals (*8–12*). Gastruloids (*10, 13–16*) are three-dimensional aggregates of pluripotent stem cells that self-organise and break symmetry via polarised Brachyury/T expression (*17*), provid-ing an ideal system to study spontaneous formation of the primary body axis *in vitro*. Recent studies suggest that cell rearrangements are the main driving mechanism of symmetry breaking in this system (*18–20*). At the same time, gene expression patterns have been shown to scale with system size, with larger gastruloids showing delayed symmetry breaking (*21–23*), suggest-ing a tight coordination between morphogenetic movements and cell fate transitions. How the interplay between these two processes drives symmetry breaking in this system remains elusive.

Here we address this question by combining cell proportion experiments with single-cell RNA sequencing, biophysical measurements, and computational modelling. By systematically varying the initial fraction of Brachyury/T-expressing cells (T+ cells), we show that the timing of symmetry breaking and subsequent gastruloid elongation depends on the initial T+ popula-tion. Mathematical modelling further reveals that cell differentiation in gastruloids is non-cell autonomous, with fate decisions emerging at the collective level. This is accompanied by a radial organisation of cell fates prior to polarisation, with T-cells preferentially localising to the aggregate periphery and T+ cells being in the core. These spatial differences correlate with surface tension variations between T+ and T-tissues, consistent with a radial cell sorting process. Finally, by integrating fate dynamics and mechanics *in silico* we are able to recapitulate the stages of symmetry breaking observed experimentally. Together, our results provide a mechanochemical framework for self-organisation in 3D embryo-like structures and establish a systematic approach to dissect cellular feedbacks *in vitro*.

## Results

### T population dynamics is dependent on the initial fraction of T+ cells in gastruloids

The ability of gastruloids to polarise and elongate is known to be dependent on the initial conditions, such as stem cell pluripotency, culture media composition or initial aggregate size (*24–27*). For example, initial heterogeneity on the levels of T or Wnt signalling are known to increase the variability in gastruloid formation (*10, 19, 25, 28, 29*). Motivated by the previous studies, we asked whether the initial state of the cells could serve as a proxy for the timing of symmetry breaking and elongation in gastruloids. To address this, we systematically varied the proportion of T+ cells at the time of seeding, following a strategy similar to the one used in chimeric embryo experiments (*30*). The canonical gastruloid protocol consists on the aggrega-tion of ↑ 300 mouse embryonic stem cells (mESC) followed by CHIR-99021 (CHIR) treatment betwen 48-72 hours post aggregation (hpa) (*10, 13, 31*). Gastruloids polarise around 72 hpa and rapidly elongate at 96 hpa. We decided to follow the same protocol but growing gastruloids in the absence of a CHIR pulse (*32, 33*) to study the inherent signalling dynamics of cells. We used an E14 Bra/T::GFP cell line to visualise T expression and cells were maintained in ES-Lif (ESLIF) medium. In these conditions, typically ↭ 5% of cells differentiate to become T+ (see Materials and Methods). We took advantage of this fact to sort T+ and T-cells from culture via fluorescence-activated cell sorting (FACS) and generate 3D aggregates with controlled T+ fractions. T+ and T-cells were mixed in different proportions to form 3D cell aggregates in ultra-low adherence multiwell plates (Fig. 1A). By using the GFP signal as a proxy for the T+ cell population, we estimated the fraction of T+ cells over time, *ω*(t) (Fig. 1B,C,D, Fig. S1A and Materials and Methods). While tissue polarisation was observed in all cases, the time of polarisation varied depending on the initial T+ fraction (Fig. 1C and Fig. S1C). The change in T dynamics depending on the inital T+ fraction became more evident when the average fraction of T+ cells (*ω*) was plotted as a function of the initial T+ fraction (*ω*(0)) at 24, 48 and 72 hpa (Fig. 1D). Cell aggregates that were initially formed with a large amount of T+ cells (*ω*(0) ↫ 0.5) polarised and elongated at 48 hpa instead of at 72 hpa (Fig. 1B,C and Fig. S1B,C). The fact that a high T+ population favors rapid polarisation is analogous to the high expression of T+ cells following the addition of CHIR in the canonical protocol, which leads to fast polarisation and elongation (*10, 13*). Notably, the timing and degree of elongation were also affected depending on the initial T+ fraction (Fig. 1E and Fig. S1B). We thus conclude that, while gastruloids can polarise and elongate irrespective of the initial fraction of T+ cells, the timing of symmetry breaking and elongation is dependent on the initial T+ fraction.

**Figure 1.**
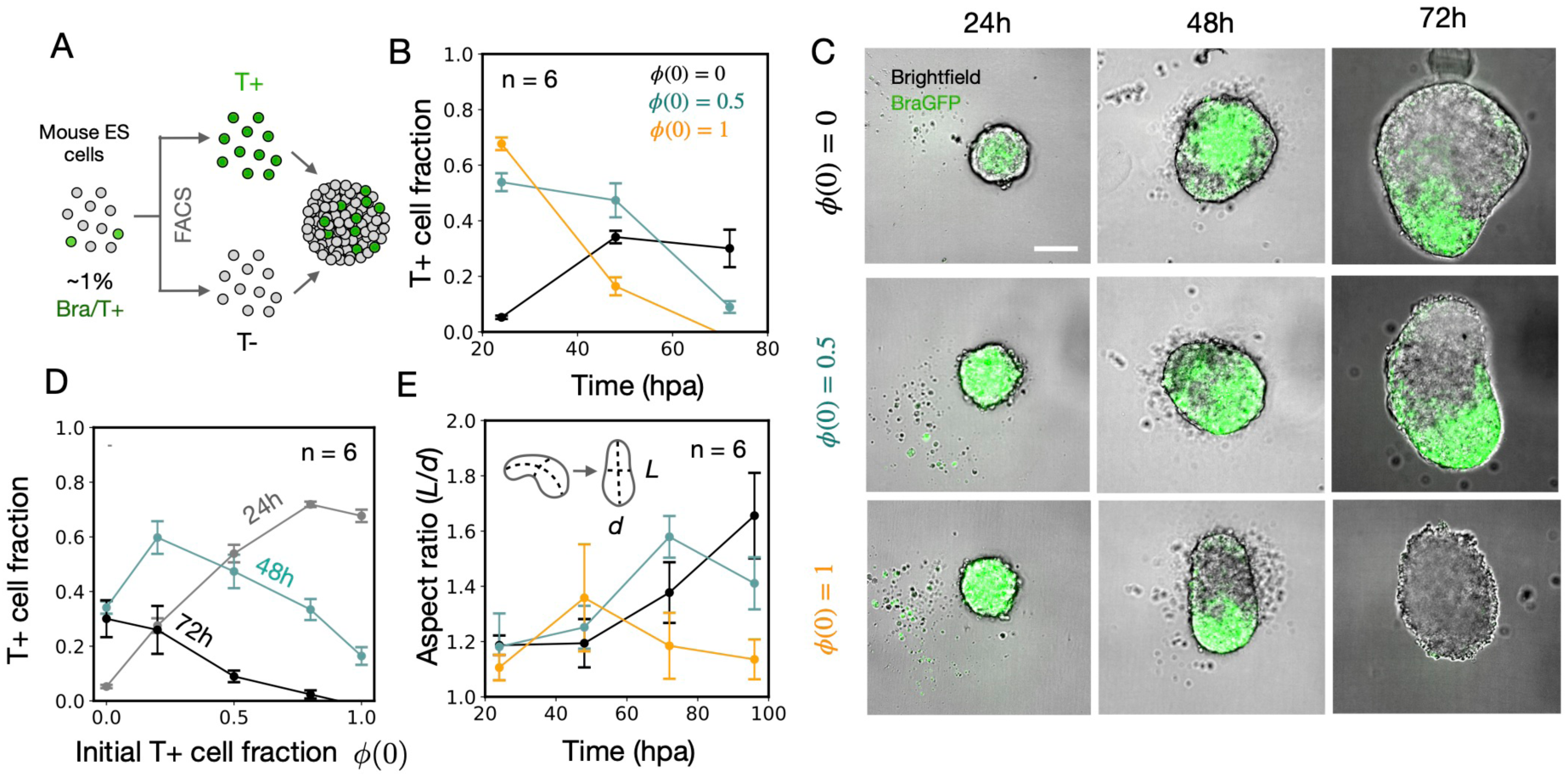
T dynamics under controlled cell proportions. A) Schematics of the cell proportion experiments. mESCs were cultured in 2D under ESLIF conditions, and were subsequently FACS sorted into T+ and T-populations. The different cell populations were mixed in different proportions to form 3D aggregates in multi-well 96 plates in N2B27 medium. B) Time evolution of the T+ fraction for different starting T+ populations *ω*(0) = 0, 0.5, 1. C) Representative brightfield images of the aggregates at 24, 48 and 72 hpa for initial T+ frac-tions *ω*(0) = 0, 0.5, 1. The images are composed with the signal from a GFP reporter of T expression. Scale bar: 200 *µ*m. D) T+ cell fraction as a function of the initial fraction of T+ cells *ω* for 24, 48 and 72 hpa. In each experiment approximately 50 cell aggregates were analysed. The fraction of T+ cells was obtained by normalising by the extrapolated GFP intensity at *ω* = 1 at 24 hpa. E) Time evolution of the aspect ratio of gastruloids for initial T+ fractions *ω*(0) = 0, 0.5, 1. Error bars in panels B and D correspond to 2 ⇑ SEM over *n* = 6 replicates. In each replicate ↑ 10 aggregates were averaged for each condition. Error bars in E indicate SD.

### Single-cell RNA-seq allows the identification of the major cellular states during early gastruloid differentiation

The cellular identity of T-cells in our experiments is compatible with both a pluripotent state as well as with a differentiated mesoderm state in which T expression was lost. In order to discern between these two cases and to allow for the identification of the major cellular states, we made use of our previously published single-cell RNA sequencing dataset (*32*). We focused on the cell fate dynamics from 24-48 hpa (prior to the addition of CHIR) by integrating both time points (Fig. 2A, gray data points in the background). Given the challenges of classical clustering and annotation approaches to split states with subtle gene modifications, we decided to use a custom annotation approach (see Materials and Methods) using cell fate markers known in the literature. Using this method, we annotated the major cellular fates and their proportions, as well as their time evolution trajectories (Fig. 2A,B, Figs. S2, S3 and Table S1). Compared to other methods which require many annotation markers (*34*), our simple method provides a robust approach more suitable for the annotation of continuous processes with small transcriptomic differences. We further validated the consistency of our annotation approach with traditional clustering methods, showing good agreement between communities discovered and annotation (see Fig. S2B, C, E). At 24 hpa, approximately one-third of the cells were found in an epiblast/pluripotent state (Nanog+, Sox2+) while the remaining two-thirds started transitioning to primitive streak-like states (see Fig. 2A and Fig. S2D). A large population of those differentiated cells (↓ 63%) were Nanog+, T+, which we may refer to as posterior epiblast-like. The remaining fraction of differentiated cells consisted mainly on a Nanog-, T+ population (↓ 31%) referred to as a posterior primitive-streak-like state and only a minority of cells (↓ 6%) were found in anterior mesoderm-like (Eomes+, Gata6+, Mesp1+) or endoderm-like (Eomes+, Gata6+, Sox17+) states (see Fig. 2A). The scenario at 48 hpa showed an increase of the posterior primitive streak-like population (Nanog-, T+) coming from cells differentiating from the posterior epiblast-like state. Finally, the differentiation of Nanog-, T+ cells contributed to the emergence of a posterior mesoderm-like state (Aldh1a2+, Tbx6+) (see Fig. 2A, C, E S2). Overall, the previous results recapitulate an analogous progression of the bifurcation tree observed in the mouse embryo (*35*). A similar analysis on a second replicate experiment resulted in the same conclusions stated above (see Fig. S3).

**Figure 2.**
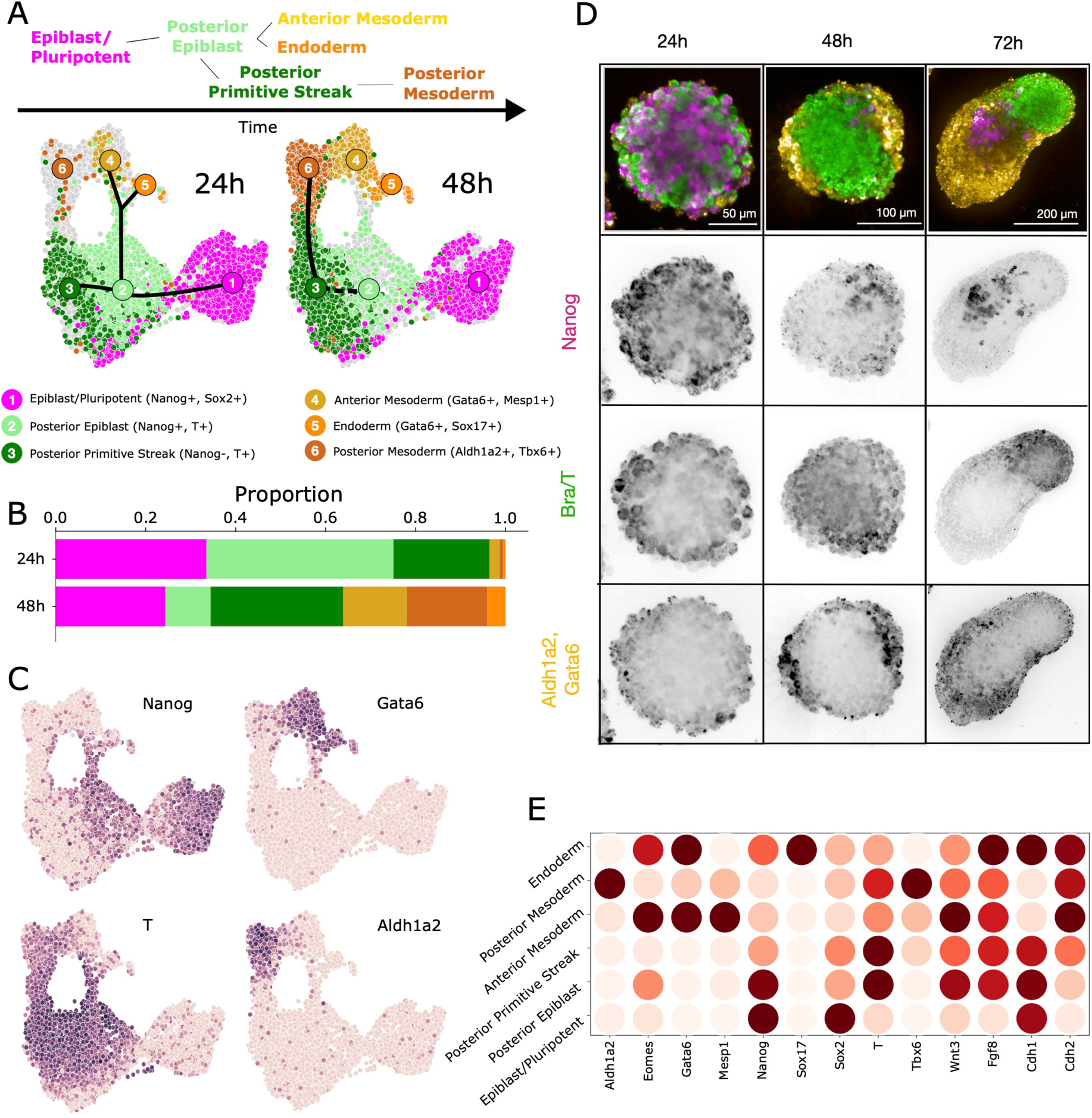
Cell fate dynamics during early gastruloid differentiation. A) Integrated UMAP of 24 hpa and 48 hpa: light gray indicates combined data and colored points correspond to individual timepoints showing how the different states are populated over time. B) The corresponding proportions by annotation are shown. C) UMAPs showing expression of selected genes. D) HCR stainings for the 3 main cellular states at 24, 48 and 72 hpa: Nanog+ pluripotent state, a T+ primitive streak-like state and further differentiated states being Aldh1a2+ and Gata6+. E) Mean normalised and log-normalised gene expression per cluster of relevant genes per annotation cluster at 48 hpa.

To verify the progression and the spatial patterning of the previously identified cellular states, we performed RNA hybridization chain reaction (HCR) stainings. Given that experimentally we have access to the dynamics of the T+ state, we broadly classify all cell types into 3 states: Nanog+ pluripotent state (state A, cluster 1), T+ primitive streak-like state (state B, clusters 2 and 3) and further differentiated states Aldh1a2+ and Gata6+ (state C, clusters 4,5 and 6). We stained for markers of these states at 24, 48 and 72 hpa in gastruloids (Fig. 2D and Fig. S4) and on purely T-(*ω* = 0) and T+ (*ω* = 1) FACS sorted aggregates (Fig. S5). Nanog was homogeneously distributed at 24 hpa for both control gastruloids and T-sorted aggregates and was found in small clusters in the anterior region at 72 hpa (see Figs. 2D,S4 and S5). Conversely, T+ aggregates showed low Nanog expression at 24 hpa and no expression at 48 hpa, with structures resembling small 72 hpa gastruloids (Fig. S5). We thus conclude that T+ cells (state B) cannot differentiate back to the pluripotent state (Nanog+, state A). Interestingly, we observed that cells in state C (expressing Aldh1a2 and/or Gata6) were restricted to the periphery in both gastruloids (Fig. 2D) and T-sorted aggregates at 48 hpa (Fig. S4, S5). Similarly, Aldh1a2 and Gata6 were preferentially found on the periphery of the 24 hpa T+ aggregates (see Fig. S5). At 72 hpa, Aldh1a2 and Gata6 were found in the anterior region of T-sorted aggregates and gastruloids, consistent with previous work (*13, 32*). Overall, our results show that radial differences in T expression are already present at 48 hpa, prior to gastruloid polarisation at 72 hpa.

### T-cells are enriched on the periphery of the 3D aggregates in cell proportion experiments

The previous HCR stainings show that state C (T-) cells form clusters on the periphery of 48 hpa gastruloids while state B (T+) cells are found on the core (Fig. 2D). Given recent evidences of cell sorting in gastruloids depending on Wnt activity (*19*), we decided to explore if state A cells (T-) would also be restricted to the periphery of the aggregates in our cell proportion experiments. Given that our cell line had a single cell reporter (Bra::GFP), it was not a priori possible to distinguish A-B from B-C transitions. In order to circumvent this problem, we made use of exogenous labelling of cells with a SiR-DNA dye (see Fig. 3A,B and Fig. S6A). When initially labelling the T-population (A state) with the dye, cells that were SiR-DNA+, T+ after 24h corresponded to T-cells that activated T (p-transition, see Fig. 3B). Conversely, when initially labelling the T+ population, cells that were SiR-DNA+, T-after 24h would correspond to T+ cells that turned off T (q-transition, see Fig. 3B). To confirm that the SiR-DNA dye was not affecting the cells, we labelled the T+ and T-populations and did not find any effect on T proportions (see Fig. S6B).

**Figure 3.**
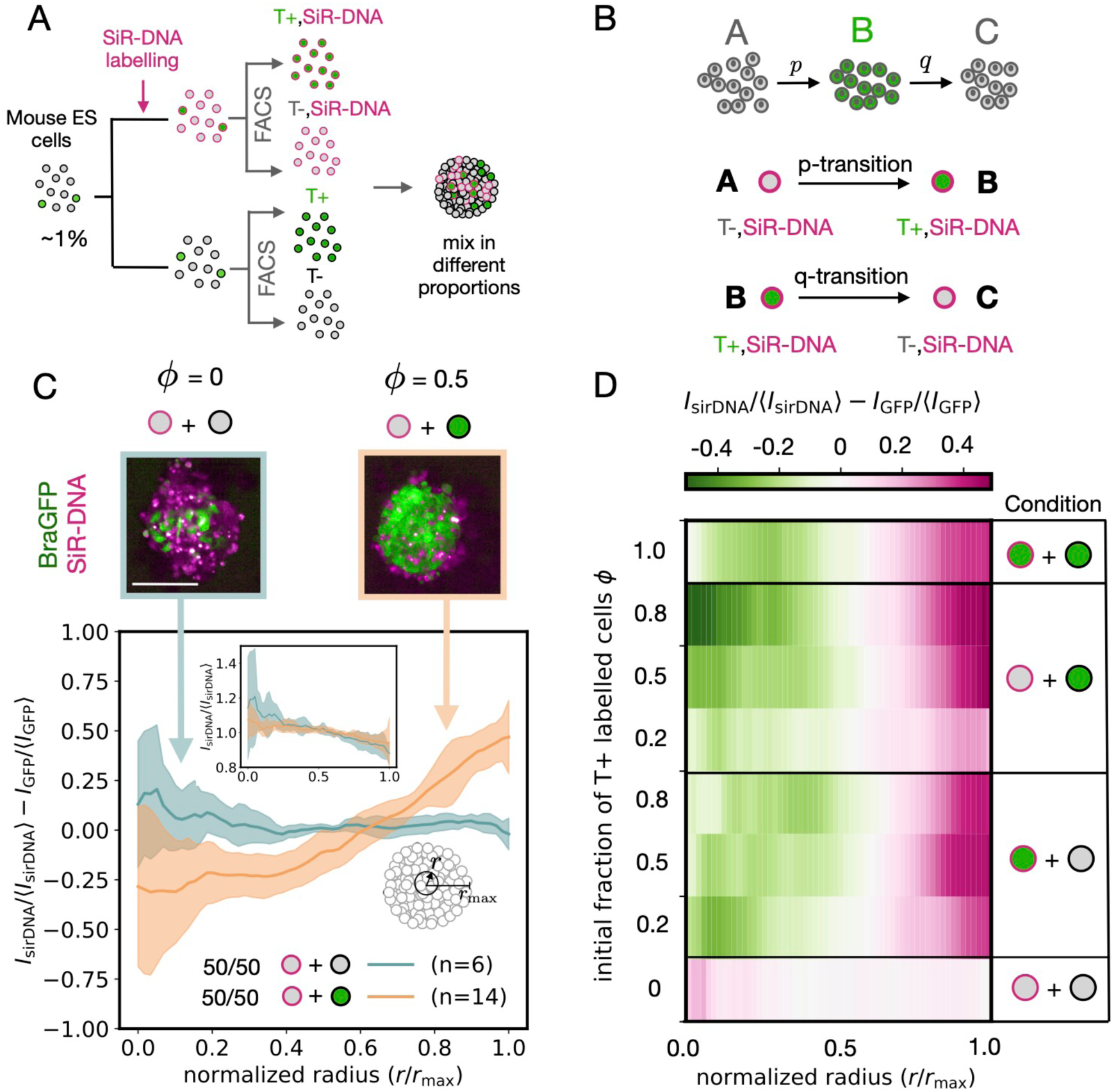
Spatial organisation of T in cell proportion experiments. A) Schematics of the cell proportions experiment with exogenous labelling. mESCs were cultured in 2D under ESLIF conditions and a certain fraction was labelled with SiR-DNA for 1h. Cells were subsequently FACS sorted into four different populations: (1) T+, SiR-DNA+; (2) T-, SiR-DNA+; (3) T+, SiR-DNA- and (4) T-, SiR-DNA-. Finally, the different cell populations were mixed in different proportions to form 3D aggregates in multi-well 96 plates in N2B27 medium and studied at 24 hpa. B) SiR-DNA labelling of the T- cells allowed us to detect *p*-transitions while labelling of T+ cells allowed us to detect *q*-transitions. C) Cell proportion experiments mixing 50/50 populations of labelled and unlabelled cells. When the cell populations are different, T-cells are enriched on the surface of the aggregate. Radial intensity profile showing GFP enrichment in the inner part of the aggregate relative to SiR-DNA signal. Inset: Radial SiR-DNA signal profile showing no significant spatial organisation. Images correspond to confocal maximum intensity projections at 24 hpa. Shaded error bars correspond to SD. D) Colormaps of the radial profile for different starting fractions of T+ cells *ω* as well as different populations labelled with SiR-DNA. In the inner part of the aggregate the majority of cells are in state B, while in the periphery the majority of cells are in either A or C states.

We first mixed 50/50 T+ and T-cells and labelled the latter population with SiR-DNA (Fig. 3C). At 24 hpa, we found that the cells that did not activate T (T-, SiR-DNA+) were enriched on the periphery of the aggregates (see Fig. 3C, upper panel). This was further quantified by computing the radial intensity profile of the difference between the normalised SiR-DNA and T signals (Fig. 3C, lower panel; see Materials and Methods). As a control, we mixed SiR-DNA labelled T- and unlabelled T- cells (50/50 ratio) and the enrichment was lost as expected (see Fig. 3C, blue case). Hence, provided that all cells are initially T-, no radial differences in the T population are found during the first 24 hpa. Interestingly, despite radial differences in T expression were found in the 50/50 mixing case, no significant spatial variations in the SiR-DNA signal were detected (Fig. 3C, inset). This result suggests that cell sorting alone cannot account for the observed spatial pattern of T expression during the first 24 hpa. However, this observation is compatible with a scenario in which cell sorting and fate transitions occur simultaneously. Indeed, if one were to assume that T- cells sort to the periphery and T+ cells sort to the core in the 50/50 mixing case, the number of labelled T- cells moving from the core to the periphery could be balanced by the number of labelled T- cells that activate at the periphery and subsequently move to the core through cell sorting. Radial differences in T expression were consistently found when mixing T+ and T- cells in different proportions (see Figs. 3D, S6 and S7). In particular, when starting with a purely T+ aggregate and labelled with SiR-DNA half of the cells, after 24h the cells that de-activated (T-, SiR-DNA+) were also enriched in the outer region while T+ cells remained enriched in the core (see Figs. S6A and S7 cases *ω* = 1). This was consistent with our HCR stainings of purely T+ aggregates at 24 hpa (see Fig. S5). Hence, our results show that T- cells are enriched on the periphery of the aggregates irrespective of their previous T expression history (i.e. both A and C states). We thus conclude that cell sorting and cell fate transitions are likely to occur simultaneously in the aggregates and contribute to the observed spatial differences in T expression. This led us to hypothesise possible differential mechanics between T+ and T- cells leading to mechanical cell sorting.

### T+ and T-cell aggregates exhibit differential tissue mechanics

To investigate if the final spatial organisation on T expression was a consequence of differential tissue mechanics, we performed fusion experiments as well as nanoindentation to characterise the tissue mechanical properties (Fig. 4A) (*23, 36, 37*). We found that pairs of T+ cell agge-gates fused completely after 10 h while pairs of T-aggregates exhibited a phenomenon known as arrested coalescence, i.e. incomplete fusion (*36*) (Fig. 4A, right and Fig. 4B). Our results align with recent findings where T KO cellular aggregates also exhibited partial fusion as opposed to wildtype aggregates (*23*). Interestingly, the dynamics of heterotypic pairs (T+,T-) was clearly different from the homotypic ones (T+,T+ or T-,T-), hinting to the presence of an interfacial tension between tissue types (Fig. 4B, orange curve). We next decided to explore if 24 hpa and 48 hpa gastruloids have similar mechanical properties compared to T- and T+ aggregates, respectively. To do this, we performed the same set of homotypic fusion experiments using 24 hpa and 48 hpa gastruloids. Indeed, similar trends were observed akin to the T- and T+ homotypic fusions, namely 24 hpa pairs exhibited arrested coalescence while 48 hpa pairs fused completely (see Fig. 4C). This is consistent with the timeline of gastruloid differentiation, where T expression is upregulated around 48 hpa. We next performed nanoindentation measurements of the aggregates to measure the effective tissue elasticity (see Materials and Methods). By combining the estimate parameters from the fusion experiments and the nanoindentation measurements, we could infer the surface tension *ε* and viscosity *ϑ* of the aggregates (*36*). We found a ↑ 3-fold increase in surface tension *ε* and a ↑2-fold increase in viscosity *ϑ* for T+ ag-gregates compared to T-aggregates (see Fig. 4D, E, F). The effective elasticity was found to be slightly lower for T+ aggregates compared to T-aggregates (Fig. 4F). While 48 hpa gastruloids showed a similar 3-fold increase in surface tension with respect to 24 hpa gastruloids, this was not the case for viscosity which remained similar. We reason that this might be a consequence of the fact that 48h gastruloids have a substantial fraction of T-cells (↑ 50 %, see Fig. 1D) that contribute to a lower tissue viscosity. Overall, our results in Fig. 4 are compatible with the classical picture of mechanical cell sorting, where the higher surface tension tissue stays in the core while the lower surface tension tissue is restricted to the periphery (*38, 39*). Well known candidate proteins driving these different tissue affinities are cadherins (*39–41*). Indeed, we confirmed such a difference at the protein level using Western blots (see Fig. S8) and found that N-cadherin (Cdh2) levels were higher in T+ aggregates than in T-aggregates. Conversely, E-cadherin (Cdh1) levels were higher in T-than in T+ aggregates. Similarly, RNA expression of the homologous cadherins showed differential expression between pluripotent and primitive streak-like states, giving additional support to this evidence (see Fig. 2E). We thus conclude that differential cadherin levels might be involved in the sorting process, in line with other studies on gastruloids (*18, 19*). In all, we find changes in the material properties of gastruloids from 24 to 48 hpa which are concomitant with the increase in T+ population during this same time period.

**Figure 4.**
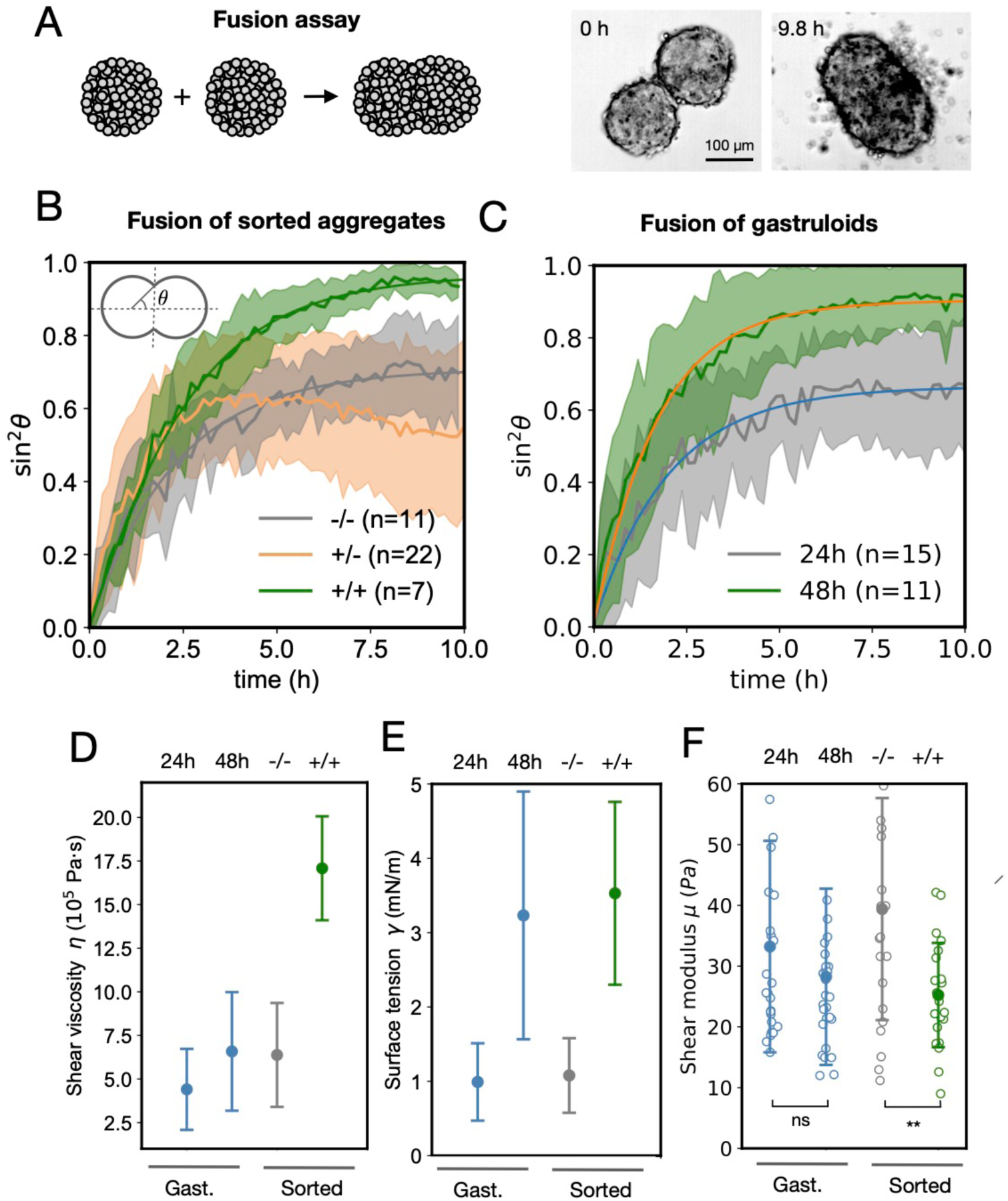
T expression impacts the mechanics of cellular aggregates. A) Fusion assay. Brightfield images of two fusing T-sorted aggregates before and after fusion. B) Time evolution of sin^2^ *ε*, where *ε* corresponds to the contact angle between the two aggregates (see inset). The average curves correspond to homotypic (T+/T+, T-/T-) and heterotypic (T+/T-) fusions of T+ and T-sorted aggregates.The average radius of T- and T+ aggregates was 80 *±* 6 *µ*m and 88 *±* 6 *µ*m, respectively (mean *±* SD). *n* corresponds the number of analysed aggregates per condition. C) Time evolution of sin^2^ *ε* for the fusion of 24 hpa and 48 hpa gastruloids. The average radius of 24 hpa and 48 hpa gastruloids was *R* = 100 *±* 7 *µ*m and *R* = 145 *±* 11 *µ*m, respectively (mean *±* SD). Solid lines correspond to fits to a mathematical model considering the aggregates behave as Kelvin-Voigt materials. D, E) Inferred effective tissue shear viscosity *ϑ* and surface tension *ϖ* for gastruloids, T+ and T-aggregates. F) Nanoindentation measurements of the shear elastic modulus *µ* for gastruloids, T+ and T-aggregates. ** *P <* 0.01, both t-test and Mann-Whitney U test. ns: both t-test and Mann-Whitney U test. The number *n* corresponds to number of fusion events. Shaded error bars correspond to SD.

### Cell-cell communication explains T proportions in the 3D cellular aggre-gates

The fact that no spatial variations in the SiR-DNA signal were found in the cell proportion experiments (Fig. 3C, inset) suggests that cell sorting is not the only mechanism at play and thus cell differentiation also plays a key role during symmetry breaking. We thus asked if cell-cell communication is required to explain T dynamics in the cell proportion experiments in Fig. 1. We first noticed that the non-monotonous behaviour at 48 hpa in Fig. 1D is incompatible with a simple linear stochastic process (*42,43*) of autonomous cell fate decisions. This is because such a process leads to a linear dependence of the fraction *ω*(t) of T+ cells at a time t on the initial fraction *ω*(0) (see Appendix). Hence, we argue that some form of cell-cell communication is necessary to account for the nonlinear dependence on the initial cell populations observed in the experiments. We considered a minimal three-state model (*43–46*) consistent with our analysis of scRNA-seq data (Fig. 5A). We performed a numerical exploration of 24 possible cell signalling models in which states A and B can control differentiation through cellular feedback (see Fig. S9 and Appendix) and found that, in the mean field approximation, the models that best fit the data correspond to the ones where the pluripotent population (state A) inhibits cell differentiation, specifically the B-C transition (see Figs. S9, S10). Such a cell-cell signalling scenario has been considered as a plausible paracrine feedback mechanism, whereby residual pluripotent cells inhibit cell differentiation (*43*). In Fig. 5A,B,C (left) and Fig. S1A we show the model that best fits our experimental data, which considers state A inhibiting both p and q-transitions. From the fit we obtain the transition rates (p = 0.14 *±* 0.01 h*^↑^*^1^, q = 0.071 *±* 0.003 h*^↑^*^1^, mean *±* SD) as well as the feedback strength parameter (K = 0.17 *±* 0.02, mean *±* SD). Hence, a simple three-state model with cell-cell communication accounts for the experimental results.

**Figure 5.**
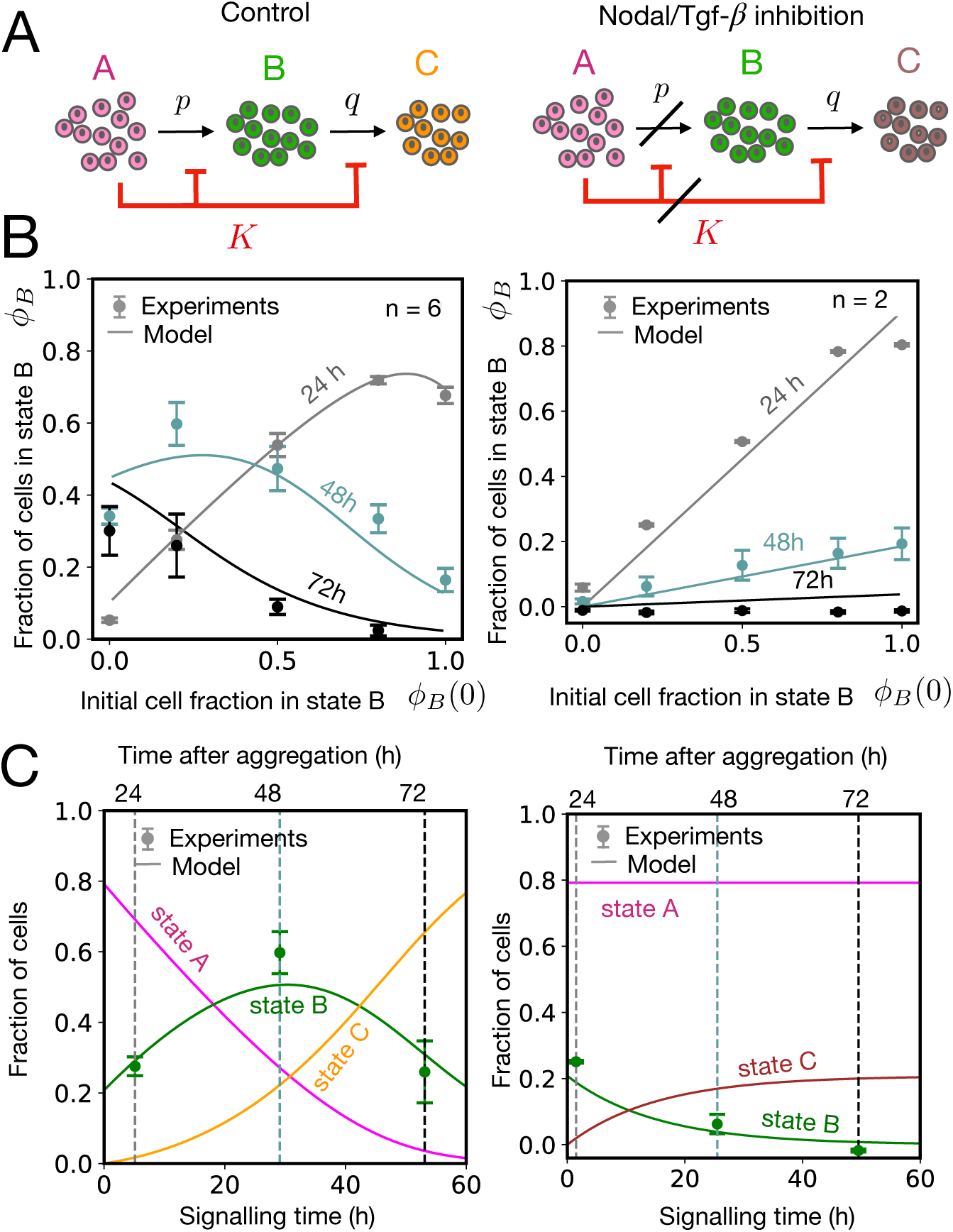
Cell-cell communication explains cell fate dynamics in gastruloids. A) Minimal cell-cell signalling model considering cells transition through 3 main cellular states (A, B and C). State A is assumed to prevent cell differentiation from A to B and from B to C with feedback strength *K*. In principle, given that the inhibitory signals might affect cell differentiation, we denote state C with a different color (brown). B) Evolution of the fraction of cells in state B as a function of the initial fraction of cells in state B for 24, 48 and 72 hpa. The fraction of cells in state B was obtained by normalising the GFP intensity by the extrapolated intensity at *ω* = 1 at 24 hpa. Solid lines correspond to the fits using the models in panel A (see Appendix). The fitted parameters for the control case (left) are *p* = 0.14 *±* 0.01 h*^→^*^1^, *q* = 0.071 *±* 0.003 h*^→^*^1^ and *K* = 0.17 *±* 0.02 (mean *±* SD). For the Nodal/Tgf-*ϱ*-inhibited case, the fit was obtained assuming *p* = 0 and *K* ⇐⇒ (no feedback) leading to *q* = 0.0660 *±* 0.0004 h*^→^*^1^. Error bars correspond to 2 ⇑ SEM over different replicates without the addition of SiR-DNA. In each replicate ↑ 10 aggregates were averaged for each condition. C) Time evolution of the three states considering *ω_A_*(0) = 0.8, *ω_B_*(0) = 0.2, *ω_C_*(0) = 0 and using the fitted parameters from the experiments (see Materials and Methods). Vertical dashed lines indicate 24, 48 and 72 hpa while the x-axis corresponds to the signalling time. The initial timepoint *t* = 0 is inferred fr_3_o_3_m the fit (see Materials and Methods).

We next asked how the parameters in our differentiation model are affected under perturbations of Wnt, Fgf and Nodal/Tgf-*β*, which are known to be the main signalling pathways involved in symmetry breaking (*13, 32, 47, 48*). In all three cases, the model showed that the p-transition was abolished, while the q-transition remained unaffected (Figs. 5, S11,S12 and S13 and Table S3). To verify the results of the model, we considered the case of 50/50 T+/T-starting populations under Fgf inhibition with PDO3. In this case, the model predicted that after 48h, the A-state cells do not transition to state B because of Fgf inhibition, while almost all B cells transition to state C. This was confirmed through HCR stainings where the T (state B) signal was almost absent while Nanog and Aldh1a2, Gata6 signals (states A and C, respectively) were the dominant ones (see Fig. S11). Our results suggest that Wnt, Fgf and Nodal/Tgf-*β* are required for primitive streak formation (state B) while leaving mesoderm formation (state C) unaffected, in line with recent findings on Wnt and Nodal signalling in gastruloids (*48*).

Finally, our model suggests that Nodal/Tgf-*β* inhibition abolishes cell-cell communication (see Fig. 5, right) as opposed to the inhibition of Wnt or Fgf signalling (see Figs. S12 and S13). This can be seen graphically in Fig. 5B, where under Nodal/Tgf-*β* inhibition the fraction of cells in state B show a linear dependence on the initial conditions, which is indicative of cell autonomous fate decisions. This is different from Wnt/Fgf inhibition, where the curves are still non-monotonous at 48 hpa (see Figs. S12 and S13). To further study the role of Nodal/Tgf-*β* signalling in cell-cell communication, we performed a mass spectrometry analysis of the molecular components of the extracellular space by collecting the supernatant of gastruloid media at 48 hpa (see Materials and Methods). Our analysis detected several proteins secreted during the first 48h by gastruloids including those involved in Nodal/Tgf-*β* pathway (Nodal and Lefty1) while proteins related to the Wnt or Fgf pathways were not detected (see Extended Data). Therefore, we propose that Nodal signalling is a key contributor to cell-cell communication during gastruloid symmetry breaking.

### Cell fate transitions and cellular rearrangements coordinate gastruloid symmetry breaking in silico

Previously we showed that the rates of cell differentiation and cell-cell rearrangements (given by the fusion rate, see Fig. 4B,C) are of the same order, suggesting that cell fate transitions and cell sorting occur at a similar timescale during symmetry breaking. In order to understand how those two processes coordinate to achieve symmetry breaking in gastruloids, we coupled our simplified three-state model with 3D agent-based computer simulations. In the simulations, cells mechanically interact with each other via effective pairwise radial forces. In addition, cell division and differentiation are incorporated as stochastic processes (see Materials and Methods and Appendix 2). For simplicity, cellular feedback is not local but mean-field in order to be able to use the fitted parameters p, q and K. The mechanical interactions between A–A, B–B, and A–B cells were chosen to qualitatively reproduce the fusion experiments in Fig. 4. The B–C interactions were selected to capture the differential affinity between the two tissues, whereas the A–C and C–C interactions were treated as free parameters. The different parameters used can be found in Tables S4, S5. We were able to recapitulate the symmetry-breaking dynamics observed *in vitro*, including cell sorting, transient core formation and polarisation (Fig. 6A). The system was initialised with a pre-defined number of cells and a certain initial fraction of A and B cells. From 24 h to 36 h, cell sorting produced a transient core of mixed A and B cells surrounded by newly differentiated C-state cells (Fig. 6A). From 60 h to 72 h, the compact B-core moved to the periphery, driving clear polarisation of the aggregate (see Movie 1). The degree of polarisation was dependent on the initial *ω* fraction (Fig. 6B, left), consistent with our experimental results (Fig. S1). Moreover, the initial size of the aggregate had a clear effect on the polarisation dynamics, with larger aggregates exhibiting a delayed symmetry-breaking (Fig. 6B, right), as reported experimentally (*22, 23*) (Fig. 6C). We conclude that a combination of cell fate dynamics and cell sorting can account for gastruloid symmetry breaking.

**Figure 6.**
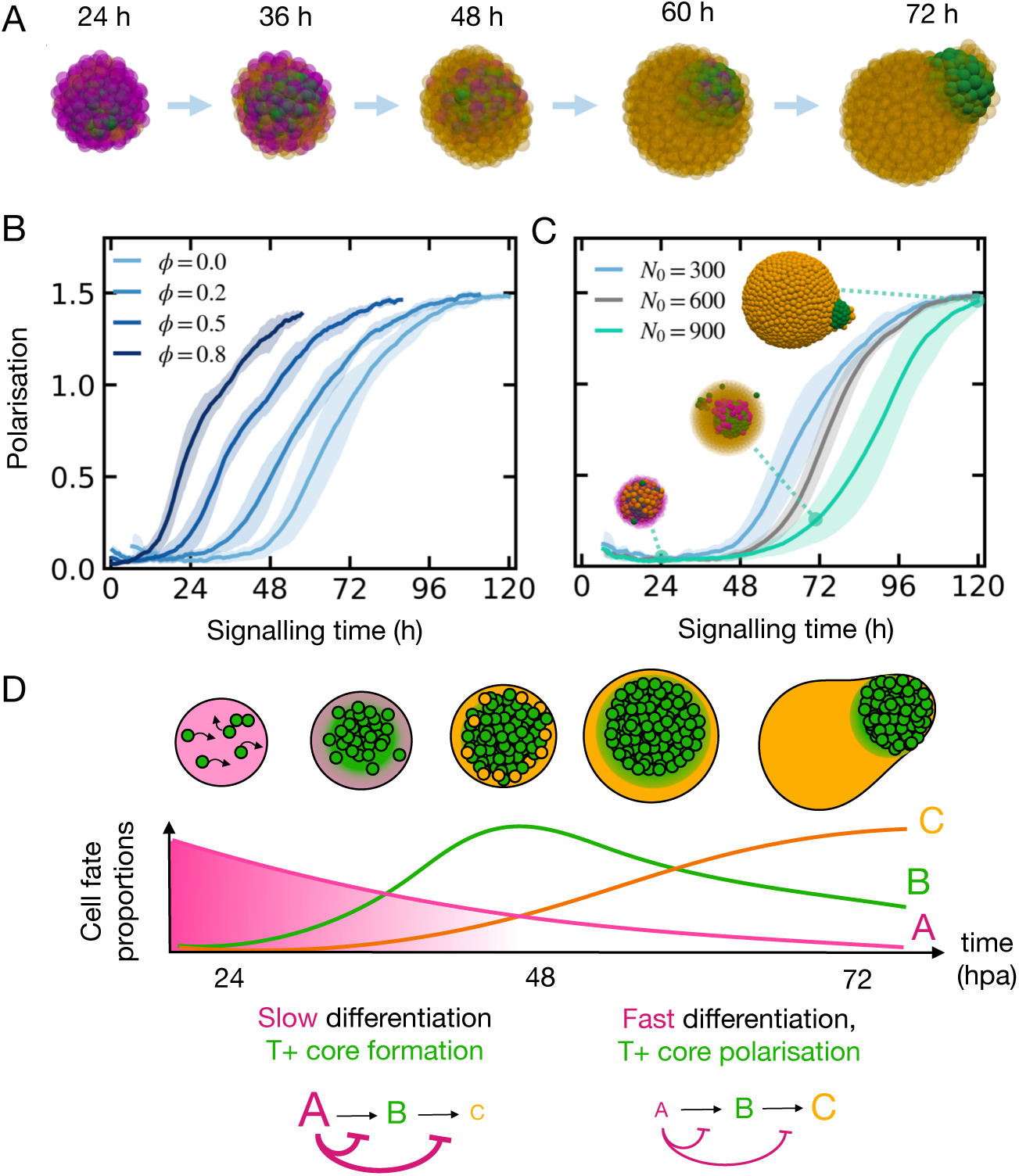
The interplay of cell fate dynamics and cell sorting explains gastruloid symmetry breaking. (A) Representative snapshots of a 3D agent-based simulation of gastruloid symmetry-breaking considering cell-cell mechanical interactions, cell proliferation and cell differentiation. Cells interact with each other depending on their cellular fate (A, B or C) adhering to one another and driving cell-cell rearrangements. (B) Polarisation vs time. Left: Initial aggregate size *N*_0_ = 300 and different initial T+ starting fractions *ω*. Right: Polarisation vs time for different initial number of cells at 0 h and initial T+ starting fraction *ω* = 0. Larger aggregates polarise slower than smaller ones. Polarisation is measured as the normalised center-of-mass distance between the B-cluster and the entire aggregate. The normalisation is done with respect to the size of the whole aggregate (see Appendix S2). For each condition, mean *±* SD is shown for 10 different simulations. (C) We propose that symmetry breaking is achieved in two stages: a first stage, where the pluripotent population slows down differentiation through cellular feedback and allows for T+ core formation through cell sorting, and a second stage where differentiation of T+ cells to the C state is sped up, leading to the polarisation process.

## Discussion

In this work, we provide a mechanochemical basis for symmetry breaking in gastruloids, taking into account cell fate transitions and collective cell rearrangements. By combining cell proportion experiments with mathematical modelling, we uncovered that cell fate decisions in gastruloids are not taken autonomously at the single-cell level, but instead emerge collectively at the population level. Notably, despite collective decision-making, we find that gastruloids’ polarisation and elongation are sensitive to the initial T+ population, in contrast to other population-level feedback mechanisms, such as in the mouse blastocyst, which buffer perturbations in initial lineage composition (*30*). This sensitivity to the initial conditions may explain the pronounced variability observed in gastruloids cultured under different medium conditions, leading to diverse cell type compositions (*26, 28, 32*). By screening a range of candidate models, we found that our experimental observations are best captured by a mechanism in which the pluripotent population slows down differentiation. Consistent with this picture, pathway perturbations and mass spectrometry analyses identify the Nodal/Tgf-*β* pathway as a likely mediator of cell-cell communication in the system. One possible hypothesis is that Lefty1 suppresses differentiation by modulating Nodal signalling in this system, analogous to its role in antero-posterior patterning in the mouse embryo (*6*). Further work will be required to directly test this hypothesis.

We further uncover spatial differences in T expression between the core and periphery of gastruloids prior to symmetry breaking. Specifically, at 48 hpa, T+ cells that subsequently lose T expression are predominantly located at the periphery of the aggregates. This transient radial organisation is reminiscent of recent work on gastruloids where primitive streak-like cells (T+) localised to the periphery while pluripotent-like cells (Sox2+) occupied the core (*29*). However, in their case the Sox2+ core polarised, leading to gastruloids with a Sox2+ pole and a T+ body (*29*), instead of the usual T+ pole. One plausible mechanism underlying this radial patterning is cell sorting driven by differential adhesion (*39, 49–53*). Consistent with this interpretation, we observe mechanical differences between T+ and T-tissues, as well as different cadherin levels, consistent with adhesion-mediated sorting. Moreover, recent experiments tracking morphogen signalling dynamics have shown that, during gastruloid polarisation, single-cell Wnt activity predicts future cell position and that axis symmetry breaking likely emerges from sorting rearrangements driven by differential adhesion (*19*). Finally, by integrating cell fate dynamics with differential adhesion-based cell sorting *in silico*, we are able to recapitulate both the transient radial organisation observed at 48 hpa and the subsequent polarisation at 72 hpa (Fig. 6), as well as the size-dependent delay in symmetry breaking reported previously (*22, 23*).

Overall, our study suggests that not only cell-cell rearrangements, but also cellular feedback coordinates symmetry breaking in gastruloids. We propose a model where symmetry breaking occurs in two different stages: a sorting-dominated stage followed by a differentiation-dominated stage (see Fig. 6D). In the first stage (24-48 hpa), the pluripotent population slows down differentiation in the system via cellular feedback, thus allowing for T+ cells to preferentially sort and form a core before extensive differentiation. In a second stage (48-72 hpa), the pluripotent population is sufficiently low such that T+ cells can differentiate significantly, creating an outer crust of mesoderm-like cells which later helps the polarisation of the T+ core via differential tissue mechanics. Hence we argue that both cell fate transitions and mechanical forces orchestrate symmetry breaking following a common morphogenetic timescale. Indeed, recent work suggests that the coupling of cell signalling and differential adhesion is necessary for robust cell sorting (*54*). Understanding how multicellular systems integrate both biochem-ical and mechanical activities to ensure robust self-organisation will help us advance in the principles of biological self-organisation.

## Materials and Methods

### Transgenic mESC cell line maintanence

Bra/T::GFP mouse embryonic stem cells (*55*) were maintained in ESLIF medium, consisting of KnockOut Dulbecco’s Modified Eagle Medium (DMEM) supplemented with 10% fetal bovine serum (FBS), 1x Non-essential aminoacids (NEEA), 50 U/mL Pen/Strep, 1x GlutaMax, 1x Sodium Pyruvate, 50 µM 2-Mercaptoethanol and leukemia inhibitory factor (LIF). Cells adhered to 0.1% gelatin-coated (Millipore, ES-006-B) tissue culture-treated 25 cm^2^ or 75 cm^2^ flasks (T25 Corning 353108; T75 Corning, 353136) in an incubator at 37 *^↓^*C and 5% CO_2_. Under these conditions, the percentage of T+ cells varied from 1 to 5% of the total population as determined by FACS.

### Cell proportion experiments

Cells were cultured in two T75 culture flasks to a confluency of ↑ 50 ↔ 80%. For the case of exogenous labelling, in the same day of FACS sorting, both flasks were washed with 15 mL PBS^+*/*+^ and one of them was incubated with 12 mL NDiff227 media (N2B27) (Takara Bio, #Y40002) containing 1.6 µM SiR-DNA (Spirochrome, SC007) for 1h while the other was incubated with N2B27 only. Four different conditions were obtained: (1) T-, SiR-DNA+, (2) T-, (3) T+, SiR-DNA+ and (4), T+ in 2 mL Eppendorf tubes containing 1.2 mL N2B27. Subsequently, cells were mixed in different proportions and seeded in 96-well U-bottom plates (Greiner Cellstar, #650970) (375 cells per well). 96-well plates of mES cell aggregates were imaged using a PerkinElmer Opera Phenix system using a x20 dry objective and N.A. 0.4 in confocal mode. A z-stack was acquired for every aggregate at 4 µm intervals.

### In situ hybidization chain reaction

Gastruloids were harvested from round bottom 96-well plates and transferred to 1.5 mL Eppendorf tubes with wide-bore p200 RNAase free tips. N2B27 media was removed after 3 washes for 5 min each with 1ml DPBS*^↑/↑^* and then samples were fixed with cold 4% PFA in DPBS*^↑/↑^* and kept at 4*^↓^*C overnight. The following day, PFA was removed from the samples by washing 3x 1mL DPBS*^↑/↑^*. At this point the samples were kept at 4*^↓^*up to a week. Next, the samples were dehydrdated and permeabilized with a series of 1 mL 100 % methanol (MeOH) washes (x4 10 min washes and x1 50 min wash at RT). The samples were rehydrated with a series of graded 1 mL MeOH/PBST (PBST: 1X PBS, 0.1% Tween 20) washes for 5 min each at RT (75/25, 50/50, 25/75, 5× 100/0). For the detection stage, a pre-hybridization step was done by adding 500 µL of probe hybridization buffer (PHB, Molecular Instruments) to the sample for 30 min at 37 *^↓^*C. The probe solution was prepared by adding 0.4 pmol of each detection probe mixture to 500 µL of PHB. The pre-hybridization solution was removed and the probe solution was added and incubated overnight at 37*^↓^*C. The next day the excess detection probes were removed by washing x4, 15 min with 500 µL probe was buffer (PWB, Molecular Instruments) which was pre-heated at 37*^↓^*C. Next, the samples were washed twice for 5 min with 5 x SSCT (5x Sodium chloride sodium citrate, 0.1% Tween in water) at RT. For the amplification stage, a pre-amplification step was done by adding 500 µL of amplification buffer (Molecular Instruments) equilibrated at RT for 30 min. Next, 6 pmol of hairpins h1 and h2 1:1 were prepared by snap cooling 2 µL from a 3 µM stock (heated at 95*^↓^*C for 90 s and cooled down to RT in the dark for 30 min). The hairpin mixture was prepared by diluting snap-cooled h1 and h2 hairpins to 1:250 in amplification buffer at RT. The amplification solution was removed from the sample and 250 µL of the hairpin mixture were added to each reaction to a final concentration of 12 nM for each hairpin. The samples were incubated at RT overnight. Finally, on the next day the excess hairpins were removed by washing with 500 µL 5x SSCT at RT (2 x 5 min, 2 x 30 min, 1 x 5 min). Samples were stored at 4*^↓^*C protected from light before microscopy. Gastruloids were transferred to flat-bottom 96 well plates (PerkinElmer Cell Carrier - 96 Ultra) and were imaged using a PerkinElmer Opera Phenix system using a x20 water and N.A. 1 in confocal mode. A z-stack was acquired for every aggregate at 4 µm intervals.

### Cell lysis and Western blots

600-cell gastruloids were pulled together at 24 hpa from a 96-well plate (↑ 6 *·* 10^4^ cells). Gastruloids were treated with 0.05 % Trypsin-EDTA (Thermo Fisch der Scientific) for 3 min. The cell suspension was washed with 5 mL DPBS^+*/*+^ and spun down for 3 min at 900 rpm twice to remove the excess Trypsin. Cells were washed twice with 5 mL ice-cold TBS-Ca^++^ buffer (5 mM CaCl_2_, 50 mM TrisHCl pH 7.5, 150 mM NaCl) containing 1 mM PMSF in DMSO and spun down for 3 min at 900 rpm. Cells were subsequently washed with 5 mL ice-cold HBSS. Finally, 250 µL RIPA lysis buffer (500 mM TrisHCl pH 7.5, 10 mM EDTA, 1.5 M NaCl, 10 % v/v Np40 and 2.5 % m/v DOC) containing one tablet of protein inhibitor cocktail diluted 1:7 was added to the cell lysate and the cells were kept on ice for 15 min. The lysate was transferred into a micro-centrifuge tube and was shaken for 1h at 48*^↓^*C. Finally, the lysate was passed through a Qia shredder and centrifuged at 15000 rpm) at 48*^↓^*C for 15 min. At this point the lysate was flash frozen and stored at ↔20*^↓^*C. To perform the Western blots, cell lysates were resolved on SDS-polyacrylamide gels (BIO-RAD Mini-Protean Precast Gels) and the proteins transferred onto nitrocellulose blotting membranes (Amersham). Membranes were blocked for 30 min at room temperature (RT) with 10 % (w/v) skimmed milk diluted in TBS (10 mM Tris-HCl [pH 7.4] and 100 mM NaCl)-0.1% Tween-20 (TBS-T), and then probed for overnight at 4*^↓^*C with primary antibodies diluted in 5% skimmed milk in TBS-T. After several washes, the membranes were incubated at RT for 45 min with horseradish peroxidase-conjugated secondary antibodies (Invitrogen) diluted in 5% skimmed milk in TBS-T. Antibody binding was detected by chemiluminiscence (ECL Prime Western Blotting Detection Reagent, Amersham) on a Fusion Fx Spectra Imager (Vilber). The antibodies used can be found in Table S2.

### Image analysis

Image analysis was done using the software MOrgAna (*56*). In order to obtain the image masks, a custom-made Python scripts for downstream data processing were used. For the cell proportion experiments, average intensities were computed at the focal plane and the background signal was subtracted. For the radial analysis, we first pre-processed the sirDNA signal I_sirDNA_ for each aggregate by removing high intensity peaks corresponding to 0.3% of the total signal. Next, the relative intensity measure *ϱ*I = I_sirDNA_/↗I_sirDNA_↘↔ I_GFP_/↗I_GFP_↘ was computed for each aggregate image, where the spatial average ↗.. .↘ was performed in the masked region. The relative intensity radial profiles for each aggregate were obtained by counting the values *ϱ*I in a set of pixels at a distance r from the center of mass of the aggregate and normalising by this value. Finally, in order to average the profiles over different aggregates, we first normalised the radial axis by interpolating the profiles. For the morphometric quantification of gastruloids, the software MOrgAna was used to straighten the gastruloids (see Ref. (*56*) for details) and the aspect ratio was defined as the ratio between the major axis L and minor axis d (see Fig. 1E and S1B). In order to quantify gastruloid polarisation, the averaged anteroposterior intensity profile I(x) was obtained by analysing several gastruloids using the software, where x was the normalised anteroposterior axis such that x ≃ [0, 1]. Finally, the polarisation parameter P was defined as 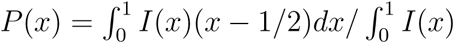.

### Nanoindentation and inference of mechanical parameters

The mechanical measurements were done using the Chiaro Nanoindenter (Optics11) adapted to a Leica DMi8 inverted microscope. The aggregates were transferred from the multiwell plates to µ-Slide 8 well coverslips (ibidi, #80826) coated with 0.1 % gelatin and containing warm N2B27. Indentations were done with a spherical cantilever probe of 27 *±* 3 µm of radius and a stiffness of 0.025 *±* 0.002 N m*^↑^*^1^. The approach speed was 5 µm s*^↑^*^1^ and the indentation depth was ↓ 6 µm (↑ 3 % of the typical size of an aggregate). The effective elastic modulus E_eff_ was calculated by fitting the Hertz’s model to the indentation curves and the corresponding shear modulus was obtained as µ = (E_eff_ /2)(1 ↔ *ς*^2^)/(1 + *ς*), assuming a Poisson’s ratio of *ς* = 1/2. The average effective elastic modulus was obtained by averaging over the different aggregates. In order to obtain the effective surface tension *ε* and viscosity *ϑ* of the cellular aggregates, the viscocapillary velocity *ε*/*ϑ* and the elastocapillary length *ε*/µ were obtained by fitting a Kelvin-Voigt model to the fusion experiments in Fig. 4B,C as explained in Refs. (*36, 37*). Finally, by using the obtained value of the shear elastic modulus µ from the nanoindentation measurements, the parameters *ϑ* and *ε* were inferred.

### Cell fate model exploration and inference of cell fate transition rates

The experimental dataset (24, 48 and 72 hpa data) was fit to the model using 4 parameters, *φ* = p/q, K, τ _24_ and τ _48_, where the last two parameters correspond to the dimensionless time-points at 24 hpa and 48 hpa. The reason why four parameters are needed instead of three (p, q, K) is because the seeding time (τ = 0) does not correspond to the time when signalling starts. Hence, an additional time parameter is needed. The timescale of the process was then obtained as 24 h/(τ_48_ - τ_24_). The dimensionless timepoint at 72 hpa was obtained as τ_72_ = 2τ_48_ - τ_24_. For the signalling perturbations, in some cases the model was simplified by considering no p-transitions or the absence of feedback (K ⇒∞) (see Table S3). The 24, 48 and 72 hpa datasets were simultaneously fit to the analytical model using non-linear least squares fitting and leave-one-out cross-validation. Parameter sets yielding a z-score larger than 1 were discarded. The final fitted parameters (see Table S3) correspond to the mean values across fits, with errors given by the standard deviation. For the model exploration shown in Figs. S9 and S10, stochastic simulations were performed to find estimations of the mean values of *ω_B_*(t) for all master equations, as closed-form solutions were not available for all 24 models. Each model was simulated with 10^4^ independent realisations and five different proportions of A and B states to compute the mean values. Parameter fitting was performed using the NOMAD algorithm (*57*), with the loss defined as the absolute error between experimental means at each time point and simulation means. The parameter search space was *φ* ≃ (0, 10),K ≃ (0, 10), τ_24_ ≃ (0, 100), τ_48_ ≃ (0, 100). The experimental data in Figs. 1 and S1A were fit to the analytical solution of the cell-cell communication model in which state A inhibits both AB and BC transitions (see Appendix Eqs. S15 and S16) using the `curve_fit` function from `scipy.optimize` in Python.

### Single-cell RNA sequencing analysis

The 24 and 48 hpa scRNA sequencing data was used from previous work in Ref. (*32*). Next, we detail the different steps of the analysis:

#### Quality control, normalisation and scaling

We analysed each sample independently corresponding to a time point each. We calculated the standard quality control metrics of total counts per cell, number of expressed genes per cell, and mitochondrial fraction. We heuristically set quality control thresholds for each metric based on the observed bimodalities of the histogram plots. Cells outside the thresholds were removed from further analysis. In the following, we used scrublet algorithm (*58*) to impute candidate doublets. We estimated the scrublet score threshold using the heuristic method described in the original paper, by observing the bimodality on the histogram of simulated doublets scrublet scores. Cells with scores above this threshold were imputed as doublets and removed from further analysis. The remaining cells in the count matrix were normalised to a total count sum of 10^5^ counts. The analysis was insensitive to the specific choice of this parameter. In addition, the matrix was log-one-plus normalised before downstream analysis.

#### Feature selection, dimensionality reduction and visualisation

For dimensionality re-duction genes were selected using the `scanpy.pp.highly_variable_genes` function with “seurat” flavor (*59*) and the top 2000 genes were used as input dimensions for a Principal Component Analysis (PCA). The results were insensitive to the specific number of top genes selected. With the PCA, we retained the 20 components with more explained variance. This choice was based on the selection of components by perfoming a randomisation test such that all datasets have the same number of components. The selection of components was also orientative and robust to the specific choice on the number of components. With the selected number of principal components, we generated a k-nearest neighbours (k-NN) graph representation of the data using the `scanpy.pp.neighbors` with correlation metric. UMAP plots were created for visualisation using the `scanpy.tl.umap` function with default parameters.

#### Annotation

For annotation, we used the marker genes appearing in Table S1. The custom annotation algorithm worked in two steps. First, cells that had at least one count of a marker gene were considered positive for the gene and negative only if there were no counts present. Each cell was assigned a fate if it expressed the combination of positive or negative markers for that specific fate. Cells that were annotated with more than one fate were treated as “Multi-assigned”, and cells with no annotation as “Unassigned”. In a second step, we trained a k-NN classifier over the PCA space with the uniquely assigned cells as the training set and the “Multi-assigned” and “Unassigned” cell as prediction cells. These two sets of cells were annotated using the majority vote of the remaining k-NN 15 nearest neighbours.

#### Time sample integration

For integrating the time samples of each dataset we combined the samples after the quality control step. With this matrix, we normalised, scaled, selected features and performed the PCA analysis in the same procedure described above. With the retained PCA components, we used the `scanpy.external.pp.harmony` function to correct for the batch effect between samples. The corrected representation was then used for constructing the graphs, the UMAPs, clustering and annotation as described before.

### Proteomics analysis

In the following section we detail the procedure used to perform the proteomic analysis of the secreted proteins in gastruloids:

#### Sample preparation

To identify proteins secreted by gastruloids, 600 cells/20 µL N2B27 media/well were seeded in a 96-well ULA plate (Greiner, 650970). For negative control the same amount of N2B27 (Takara, Y40002) media was kept. At 48 hpa, samples were collected as bulk and centrifuged at 900 rpm for 3 min. The supernatant and the control media were then filtered by Amicon Ultra-0.5 centrifugal filter unit (Merck, UFC500308) until volume was reduced to half. Three technical replicas (both for supernatant and the control media, 40 µL/well) were loaded onto separate SDS gels and stained with Coomassie blue. Gel pieces were excised (excluding BSA and proteins of similar size (55-75 kDa)) and proteins were subjected to in-gel digestion with trypsin. Next, gels band were destained with 100 mM NH_4_HCO_3_ in 40 % acetonitrile, reduced with dithiothreitol (2 µmol, 30 min, 56 *^↓^*C), alkylated in the dark with iodoacetamide (11 µmol, 30 min, 25 *^↓^*C), dehydrated with acetonitrile for trypsin digestion (400 ng, 37 *^↓^*C, 8h, Promega cat # V5113).After digestion, peptide mix was acidified with formic acid prior to LC-MS/MS analysis and desalted with a MicroSpin C18 column (The Nest Group, Inc) prior to LC-MS/MS analysis.

#### Chromatographic and mass spectrometric analysis

Samples were analysed using a Orbitrap Fusion Lumos mass spectrometer (Thermo Fisher Scientific, San Jose, CA, USA) coupled to an EASY-nLC 1200 (Thermo Fisher Scientific (Proxeon), Odense, Denmark). Peptides were loaded directly onto the analytical column and were separated by reversed-phase chromatography using a 50-cm column with an inner diameter of 75 µm, packed with 2 µm C18 particles (Thermo Fisher Scientific, cat # ES903). Chromatographic gradients started at 95% buffer A and 5% buffer B with a flow rate of 300 nl/min and gradually increased to 25% buffer B and 75% A in 79 min and then to 40% buffer B and 60% A in 11 min. After each analysis, the column was washed for 10 min with 100% buffer B. Buffer A: 0.1% formic acid in water. Buffer B: 0.1% formic acid in 80% acetonitrile. The mass spectrometer was operated in positive ionisation mode with nanospray voltage set at 2.4 kV and source temperature at 305*^↓^*C. The acquisition was performed in data-dependent acquisition (DDA) mode and full MS scans with 1 micro scans at resolution of 120,000 were used over a mass range of m/z 350-1400 with detection in the Orbitrap mass analyser. Auto gain control (AGC) was set to ‘standard’ and injection time to ‘auto’. In each cycle of data-dependent acquisition analysis, following each survey scan, the most intense ions above a threshold ion count of 10000 were selected for fragmentation. The number of selected precursor ions for fragmentation was determined by the “Top Speed” acquisition algorithm and a dynamic exclusion of 60 seconds. Fragment ion spectra were produced via high-energy collision dissociation (HCD) at normalised collision energy of 28 % and they were acquired in the ion trap mass analyser. AGC and injection time were set to ‘Standard’ and ‘Dynamic’, respectively and isolation window of 1.4 m/z was used. Digested bovine serum albumin (New England biolabs cat # P8108S) was analysed between each sample to avoid sample carryover and to assure stability of the instrument and QCloud has been used to control instrument longitudinal performance during the project.

#### Data analysis

Acquired spectra were analysed using the Proteome Discoverer software suite (v2.5, Thermo Fisher Scientific) and the Mascot search engine (v2.6, Matrix Science). The data were searches against a Swiss-Prot mouse database (as in March 2021, 17238 entries) plus a list of common contaminants and all the corresponding decoy entries. For peptide identification a precursor ion mass tolerance of 7 ppm was used for MS1 level, trypsin was chosen as enzyme and up to three missed cleavages were allowed. The fragment ion mass tolerance was set to 0.5 Da for MS2 spectra. Oxidation of methionine and N-terminal protein acetylation were used as variable modifications whereas carbamidomethylation on cysteines was set as a fixed modification. False discovery rate (FDR) in peptide identification was set to a maximum of 1%.

### Agent-based simulations

Simulations were performed using the Julia package `CellBasedModels.jl` (https://github.com/dsb-lab/CellBasedModels.jl). A center-based model was used to describe the dynamics of each cell in the aggregate taking into account repulsion and adhesion forces, as well as friction forces between cells (*60*) (Appendix S2). In addition, cells proliferated and differentiated according to the three-state differentiation model described in the Main Text (see Appendix S1). Polarisation was quantified as the normalised distance between the center of mass of the largest T+ (B-type) cluster and that of the entire aggregate. Simulations were run for times up to 96 h signalling time (the time elapsed since the onset of cell differentiation) covering the time window of the *in vitro* experiments. Starting from an initial aggregate size of N_0_ = 300 cells we simulated cases with initial fractions of T+ cells of *ω* = 0, 0.2, 0.5 and 0.8. To examine the effect of the initial aggregate size, additional simulations were performed with initial aggregate sizes of N = 300, 600, and 900 cells and *ω* = 0. For each case, 10 independent simulations were run to ensure statistical robustness.

## Code availability

All scRNA-seq analysis was performed in Python. The scripts can be found in GitHub. To maintain reproducibility, we performed the analysis using a docker image containing all the required software to reproduce the analysis, provided in Dockerhub, and using the v0.6 image. Fitting of the mean field model was performed using Julia. The scripts can be found in GitHub. Regarding the Python code to analyse the data corresponding to the cell proportion experiments, gastruloid morphogenesis, fusion experiments and nanoindentation, it can be found in GitHub.

## Acknowledgments

We thank S. Gsell, P.-F. Lenne, A. Martinez Arias, V. Ruprecht, J. Sharpe, K. Stapornwongkul and V. Weichselberger for critical reading of the manuscript. We thank the Mesoscopic Imaging Facility (EMBL Barcelona) for assistance with imaging and the Tissue Engineering Unit (CRG) for culture media preparation. The proteomics analyses were performed in the CRG/UPF Proteomics Unit which is part of the Spanish National Research Infrastructure ICTS OmicsTech. D.O. acknowledges funding from Juan de la Cierva Incorporación with Project no. IJC2018-035298-I from the Spanish Ministry of Science, Innovation and Universities (MCIU/AEI). D.O, S.S., J.G.-O., and V.T acknowledge funding by the Spanish Ministry of Science and Innovation and FEDER, under projects FIS2017-92551-EXP, PID2021-128269NA-I00, PID2024-160263NB-I00, PID2024-155925NA-I00, by the “Maria de Maeztu” Program for Units of Excellence in R&D (grant CEX2024-001431-M), and by the Generalitat de Catalunya (ICREA Academia program). G.T-C. was supported by an FPU doctoral fellowship from the Spanish Ministry of Education and Universities (reference FPU18/05091) and an EIPOD4 fellowship under the Marie Skłodowska-Curie Actions COFUND 4 (847543). S.S. acknowledges funding from the ARISE2 program funded by the European Union’s Horizon Europe research and innovation programme under the Marie Skłodowska-Curie grant agreement with No 101178241. This work was supported by funds from EMBL and ERC-2022-SYG (101072123) to V.T, and ERC-2024-SYG (101167121) to J.G.O. We thank all the members of the Trivedi group for feedback on the project.

## Author contributions

Conceived and designed the experiments: D.O., V.T. Performed the experiments: D.O., Kr.A., D.F.-M., E.H., Ke.A. Analyzed the data: D.O., G.T.-C., S.S, J.G.-O. Contributed materials/analysis tools: D.O., G.T.-C., S.S, J.G.-O., V.T. Wrote the paper: D.O., G.T.-C., S.S., J.G.-O., V.T.

## Notes

### Competing Interest Statement

The authors have declared no competing interest.

### Summary of Updates

Fig. 6 has been revised after peer review.

