## Supplementary Material for "Collective fate decisions and cell rearrangements underlie gastruloid symmetry breaking"

### S1 Three-state model for cell fate dynamics

Here we use a heuristic approach considering cells transition between different states (*43–46*). We will simplify the branched lineage process obtained using single-cell RNA sequencing analysis in Fig. 2 by combining states 2 and 3 into a single state (B) and states 4, 5 and 6 into a single state (C). Therefore, we will assume unbranched unidirectional transitions between states A (state 1), B and C. Next, we describe the different models discussed in the main text.

#### Autonomous cell fate dynamics

In the following section, we define the fraction of cells in each state as $\phi_i$, $i = A, B, C$. The simplest possible model is to assume each cell evolves through those states with rates that are unaffected by the presence of other cells. Such a model reads:

$$\dot{\phi}_A = -p\phi_A \tag{S1}$$

$$\dot{\phi}_B = p\phi_A - q\phi_B \tag{S2}$$

$$\dot{\phi}_C = q\phi_B \tag{S3}$$

where $\phi_A + \phi_B + \phi_C = 1$. For the case $p \neq q$, the solution to this linear model reads:

$$\phi_A(t) = \phi_A^0 e^{-pt} \tag{S4}$$

$$\phi_B(t) = (1 - \phi_A^0)e^{-qt} + \frac{\phi_A^0 p}{q - p}\left(e^{-pt} - e^{-qt}\right) \tag{S5}$$

where $\phi_A^0 \equiv \phi_A(0)$ and $\phi_C(t) = 1 - \phi_B(t) - \phi_A(t)$. For the particular case $p = q$, the solution takes the form:

$$\phi_A(t) = \phi_A^0 e^{-pt} \tag{S6}$$

$$\phi_B(t) = e^{-pt}\left[1 + \phi_A^0(pt - 1)\right] \tag{S7}$$

This extremely simple model can successfully reproduce a maximum in the T+ population over time; however, due to its linear nature, it cannot reproduce the nonlinear dependence on $\phi$ observed experimentally in Fig. 1D.

#### Collective cell fate dynamics

Next we consider the case of cell-cell signalling in the cellular aggregate. Let us define the transition rates $\Gamma_{AB}^{(i,\mu)}(\phi_i)$, $\Gamma_{BC}^{(i,\mu)}(\phi_i)$, where $\mu$ indicates the type of feedback that applies to the transition ($\mu = +1$, positive; $\mu = 0$, no feedback; $\mu = -1$, negative) and $i$ is the cell population that drives the feedback ($i = A, B$). For simplicity, we will exclude the models where two different cell populations feedback to the same transition (e.g. state A and B, driving a positive feedback on transition AB). A given model is defined by the type of feedback for transitions AB and BC ($\mu$ and $\nu$, respectively) and the cell population that controls such feedbacks ($i$ and $j$, respectively):

$$\dot{\phi}_A = -\Gamma_{AB}^{(i,\mu)}(\phi_i)\phi_A \tag{S8}$$

$$\dot{\phi}_B = \Gamma_{AB}^{(i,\mu)}(\phi_i)\phi_A - \Gamma_{BC}^{(j,\nu)}(\phi_j)\phi_B \tag{S9}$$

$$\dot{\phi}_C = \Gamma_{BC}^{(j,\nu)}(\phi_j)\phi_B \tag{S10}$$

where $\Gamma_{AB}^{(i,\mu)}(\phi_i) = p\Gamma^{(i,\mu)}(\phi_i)$, $\Gamma_{BC}^{(j,\nu)}(\phi_j) = q\Gamma^{(j,\nu)}(\phi_j)$ and $\Gamma$ is a modulation of the transition rate according to the feedback taking the form:

$$\Gamma^{(i,\mu)}(\phi_i) = \delta_{\mu,0} + \frac{\phi_i\delta_{\mu,1} + K\delta_{\mu,-1}}{K + \phi_i} \tag{S11}$$

where $\delta_{ij}$ corresponds to a Kronecker delta. We can rewrite the equations in a dimensionless form by rescaling time by one of the transition rates (we will use $q$ for convenience) such that we obtain:

$$\dot{\phi}_A = -\alpha\Gamma^{(i,\mu)}(\phi_i)\phi_A \tag{S12}$$

$$\dot{\phi}_B = \alpha\Gamma^{(i,\mu)}(\phi_i)\phi_A - \Gamma^{(j,\nu)}(\phi_j)\phi_B \tag{S13}$$

$$\dot{\phi}_C = \Gamma^{(j,\nu)}(\phi_j)\phi_B \tag{S14}$$

where $\alpha = p/q$ and we have the constraint $\phi_C(t) = 1 - \phi_B(t) - \phi_A(t)$. For the particular case $\mu = -1$, $\nu = -1$ and $i = j = A$, it is possible to obtain a closed-form expression for the cell fractions given that the dynamics of A is uncoupled from the dynamics of B and C. The set of equations in this case reads:

$$\begin{aligned}\dot{\phi}_A &= -\frac{\alpha\phi_A}{1+\phi_A/K}\\ \dot{\phi}_B &= \frac{\alpha\phi_A - \phi_B}{1+\phi_A/K}\end{aligned} \tag{S15}$$

and the solution reads:

$$\begin{aligned}\phi_A(t) &= KW\left(\frac{\phi_0}{K}e^{-\alpha t+\phi_0/K}\right)\\ \phi_B(t) &= \frac{1}{1-\alpha}\left[\alpha\phi_A(t) + (1-\phi_0-\alpha)\left(\frac{\phi_A(t)}{\phi_0}\right)^{1/\alpha}\right], \quad \alpha \neq 1, \alpha \neq 0\\ \phi_B(t) &= (1-\phi_0)\exp\left(-\frac{t}{1+\phi_0/K}\right), \quad \alpha = 0\\ \phi_B(t) &= \frac{\phi_A(t)}{\phi_0}\left[1-\phi_0+\phi_0\log\left(\frac{\phi_0}{K}\right) - \phi_0\log\left(\frac{\phi_A(t)}{K}\right)\right], \quad \alpha = 1\end{aligned} \tag{S16}$$

where $\phi_0 \equiv \phi_A(0) = 1 - \phi_B(0)$ and $W$ is the Lambert W function.

### S2 Agent-based model details

We next describe the details of the agent-based model used in the main text which combines cell-cell mechanical interactions and the previously described non-autonomous cell fate dynamics. The implementation is done in Julia using the package `CellBasedModels.jl`

#### Cell Mechanics

We consider the overdamped limit where each cell $i$ at position $\mathbf{x}_i$ follows the dynamics (*60,61*):

$$\lambda_s \mathbf{v}_i + \lambda_r \sum_j (\mathbf{v}_i - \mathbf{v}_j) = \sum_j \mathbf{F}_{ij} \tag{S17}$$

where $\mathbf{v}_i = \dot{\mathbf{x}}_i$, $j$ runs over the nearest neighbours and friction forces on the left-hand side of the equation balance the cell-cell interaction forces on the right-hand side. The parameter $\lambda_s$ corresponds to an effective cell-ECM friction while $\lambda_r$ describes the relative cell-cell friction. On the other hand, $\mathbf{F}_{ij}$ corresponds to the cell-cell forces. Eq. S17 can be rewritten as:

$$\left(\frac{\lambda_s}{\lambda_r n_i} + 1\right)\mathbf{v}_i = \frac{1}{n_i \lambda_r} \sum_j \mathbf{F}_{ij} + \frac{1}{n_i} \sum_j \mathbf{v}_j \tag{S18}$$

where $n_i$ is the number of nearest neighbours of the $i-$th cell. In our 3D aggregates, we expect cell-cell friction forces to dominate over cell-ECM forces i.e. $\lambda_s \ll \lambda_r$. Using the latter assumption we define a small parameter $\epsilon \equiv \lambda_s/(\lambda_r n_i) \ll 1$ and rewrite force balance as:

$$\mathbf{v}_i = \frac{1}{\lambda n_i} \sum_j \mathbf{F}_{ij} + \frac{1}{n_i} \sum_j \mathbf{v}_j \alpha. \tag{S19}$$

where $\alpha \equiv (\epsilon + 1)^{-1} \lesssim 1$ and $\lambda \equiv \frac{\lambda_r}{\alpha}$ is a friction coefficient. For numerical stability $\epsilon$ needs to be non-zero (or equivalently $\alpha < 1$). For simplicity, in our simulations we set $\alpha$ to a constant value slightly smaller than 1.

**Cellular forces**: $\mathbf{F}_{ij}$ is an effective force that accounts for cell-cell adhesion, volume exclusion and long-ranged cell-cell attractive interactions. The force depends on the position of the two interacting cells $(i, j)$ as well as on their states $(s_i, s_j)$:

$$\mathbf{F}_{ij}(s_i, s_j, d_{ij}) = f(d_{ij})\,\hat{\mathbf{r}}_{ij} \cdot \begin{cases} F_{\text{rep}}^{(s_i,s_j)}, & \text{if } d_{ij} \leq 2r \\ F_{\text{att}}^{(s_i,s_j)}, & \text{if } 2r < d_{ij} < 2\mu_{ij} r \\ 0, & \text{otherwise} \end{cases} \tag{S20}$$

where $s_i, s_j$ correspond to the state ($A, B$ or $C$) of the $i-$th and $j-$th cell, respectively; $\hat{\mathbf{r}}_{ij} = \frac{(\mathbf{x}_i - \mathbf{x}_j)}{d_{ij}}$ is the unit vector of the force, $d_{ij} = |\mathbf{x}_i - \mathbf{x}_j|$ is the distance between the two cells and $f(d_{ij}) = \left(\frac{2r}{d_{ij}} - 1\right)\left(\frac{2\mu_{ij} r}{d_{ij}} - 1\right)$. The parameters $F_{\text{rep}}^{(s_i,s_j)}, F_{\text{att}}^{(s_i,s_j)}$ set the repulsive and attractive force scale and depend on the cellular state $(s_i, s_j)$, thus incorporating homotypic and heterotypic interactions. Finally, the parameter $r$ corresponds to the typical cell radius while $\mu_{ij}$ is used to set the range of the force interaction between the two cells.

### Cell divisions

Cells divide according to the following rules: 1) A division axis is randomly chosen over the unit sphere. 2) The two daughter cells are generated and positioned on opposite sides of the division axis, centred at the mother cell's location. The centres of mass of the daughter cells are placed at a distance equal to $0.3r$, after which the mother cell is removed. 3) Each daughter cell is assigned a new division time, independently sampled from a uniform distribution with mean $\tau_{div}$ and standard deviation $\sigma_{\text{div}}$ (*30*):

$$\text{U}(\tau_{\text{div}}(1 - \sigma_{\text{div}}), \tau_{\text{div}}(1 + \sigma_{\text{div}})) \tag{S21}$$

To prevent artificially large velocities caused by strong repulsive forces immediately after cell division, each daughter cell is assigned a relaxation time $\tau_{\text{rel}}$. During this period, we assume that the daughter cells do not contribute to their neighbours' relative friction in Eq. (S17). We estimate the relaxation time to be roughly inversely proportional to the number of interactions with neighbouring cells. Therefore, we choose that when the cumulative number of interactions reaches a certain value $N_{\text{rel}}$, the daughter cell has relaxed.

#### Non-dimensional variables and parameters

By non-dimensionalising equation (S19) with respect to a typical distance $R_0$, force $F_0$ and timescale $T_0$, we obtain the equation of motion that is used in the simulations for each cell:

$$\tilde{\mathbf{v}}_i = \frac{1}{\tilde{\lambda} n_i} \sum_j \tilde{\mathbf{F}}_{ij} + \frac{1}{n_i} \sum_j \tilde{\mathbf{v}}_j \alpha \tag{S22}$$

where $\tilde{\lambda} \equiv \lambda R_0/(F_0 T_0)$, $\tilde{v}_i \equiv v_i T_0/R_0$ and $\tilde{F}_{ij} \equiv F_{ij}/F_0$. A similar approach is carried out in order to define all the dimensionless parameters in the simulations, which are denoted by a tilde. $R_0 = 5$ $\mu$m is taken as the typical cell radius in gastruloids (*62*), the typical timescale is set as $T_0 = R_0/V_0 = 4$ min, where $V_0 \simeq 1.25$ $\mu$m/min is taken as the typical cell instantaneous speed in gastruloids (*26*), and $F_0 = 100$ pN is chosen as the typical cell-cell interaction force.

#### Polarisation

The degree of polarisation is calculated at every time-step as the normalised distance between the centre of mass of the largest (T+) B-cluster and the centre of mass of the whole aggregate. We use a minimum cluster size of at least $10\%$ of the number of cells of the initial aggregate size in order for it to be considered as such and a cutoff distance of $2r$ for a cell to be considered part of the cluster. The distance between the two centres of mass is then normalised by the size of the whole aggregate. To estimate the size of the aggregate we use the radius of gyration (the root mean square distance of the constituent cells from the cluster centre of mass):

$$R_g = \sqrt{\frac{1}{N} \sum_{i=1}^{N} (\mathbf{x}_i - \mathbf{x}_{\text{CM}})^2} \tag{S23}$$

where $\mathbf{x}_i$ are the positions of the cells in aggregate of size $N$ and $\mathbf{x}_{\mathrm{CM}}$ is the position of the aggregate's centre of mass.

### Supplementary Figures

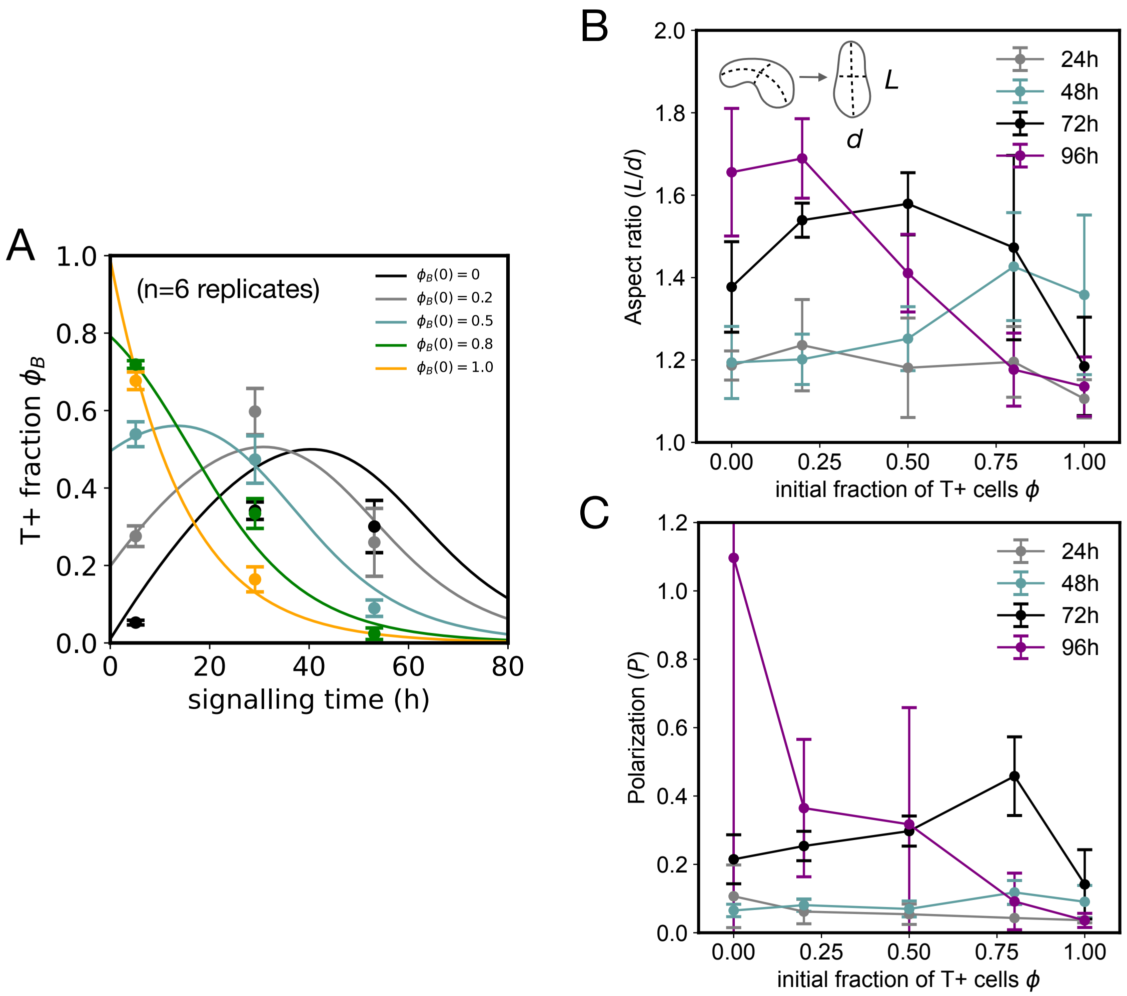


Figure S1: **Initial T+ fraction $\phi$ controls the timing of gastruloid polarisation and elongation**. A) Time evolution of the T+ fraction for different initial conditions. The data is the same as in Fig. 1B,C with the solid lines indicating the corresponding fits to the model. Mean $\pm$ 2 $\times$ SEM. $n = 6$ replicates. B,C) Polarisation and elongation of gastruloids for different initial T+ fractions $\phi$. The polarisation $P$ was calculated as the first dipolar moment (see Materials and Methods). Mean $\pm$ SD, $n = 5$ replicates.

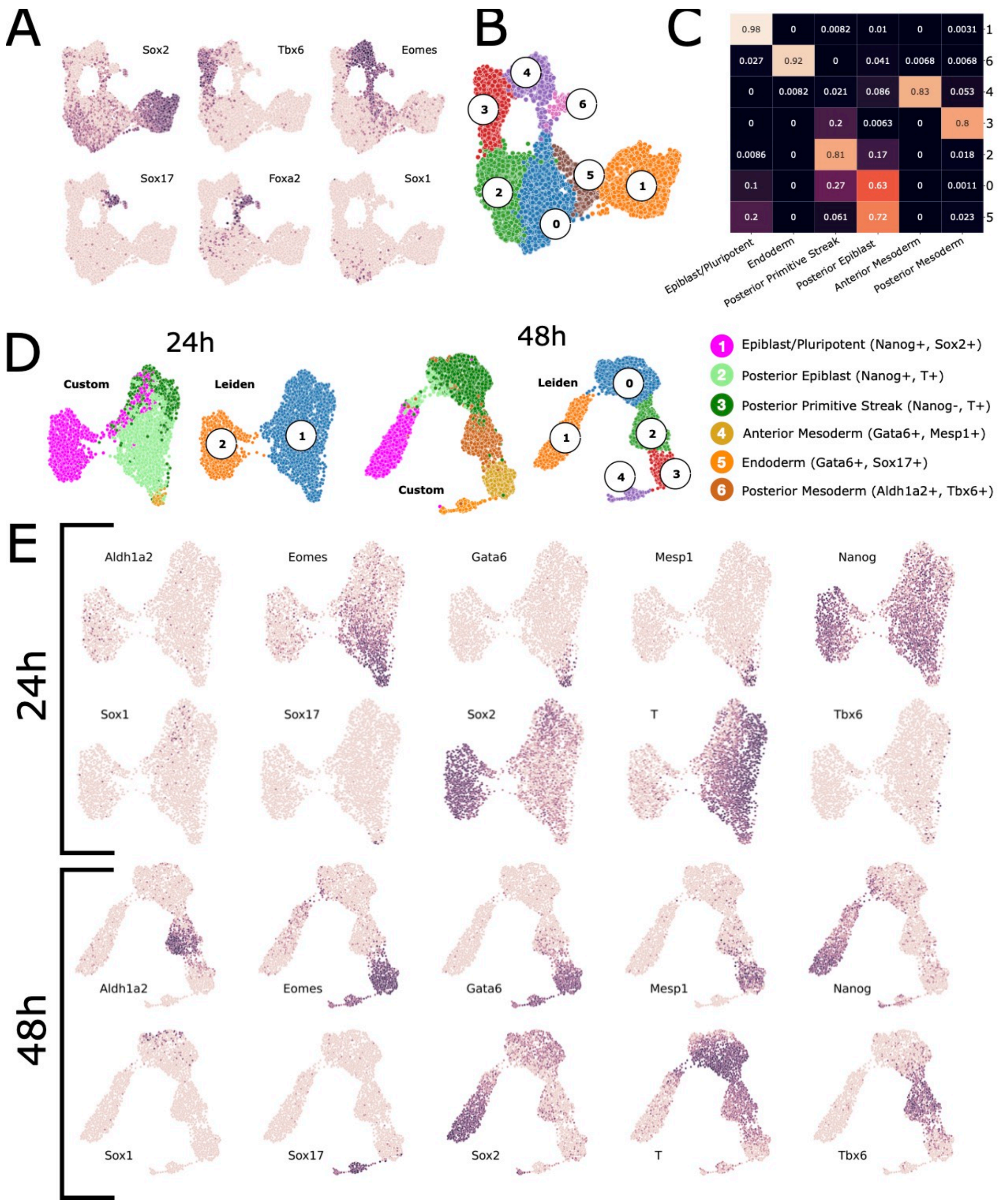


Figure S2: **Additional information of the scRNA-seq dataset presented in the main text**. A) UMAPs integrated analysis (24 hpa and 48 hpa) colored by marker genes used for the annotation of the clusters, in addition to the markers shown in Fig. 2. B) UMAP integrated analysis (24 hpa and 48 hpa) with the identified leiden clusters. C) Confusion matrix normalised by columns showing the consistency of annotation between the annotated clusters in the independent analysis and the integrated analysis cluster annotations. D) UMAPs of the independent analysis at 24 hpa and 48 hpa of the dataset, with the obtained clusters annotated. E) Marker genes expressed over the UMAPs of the time-independent analyses.

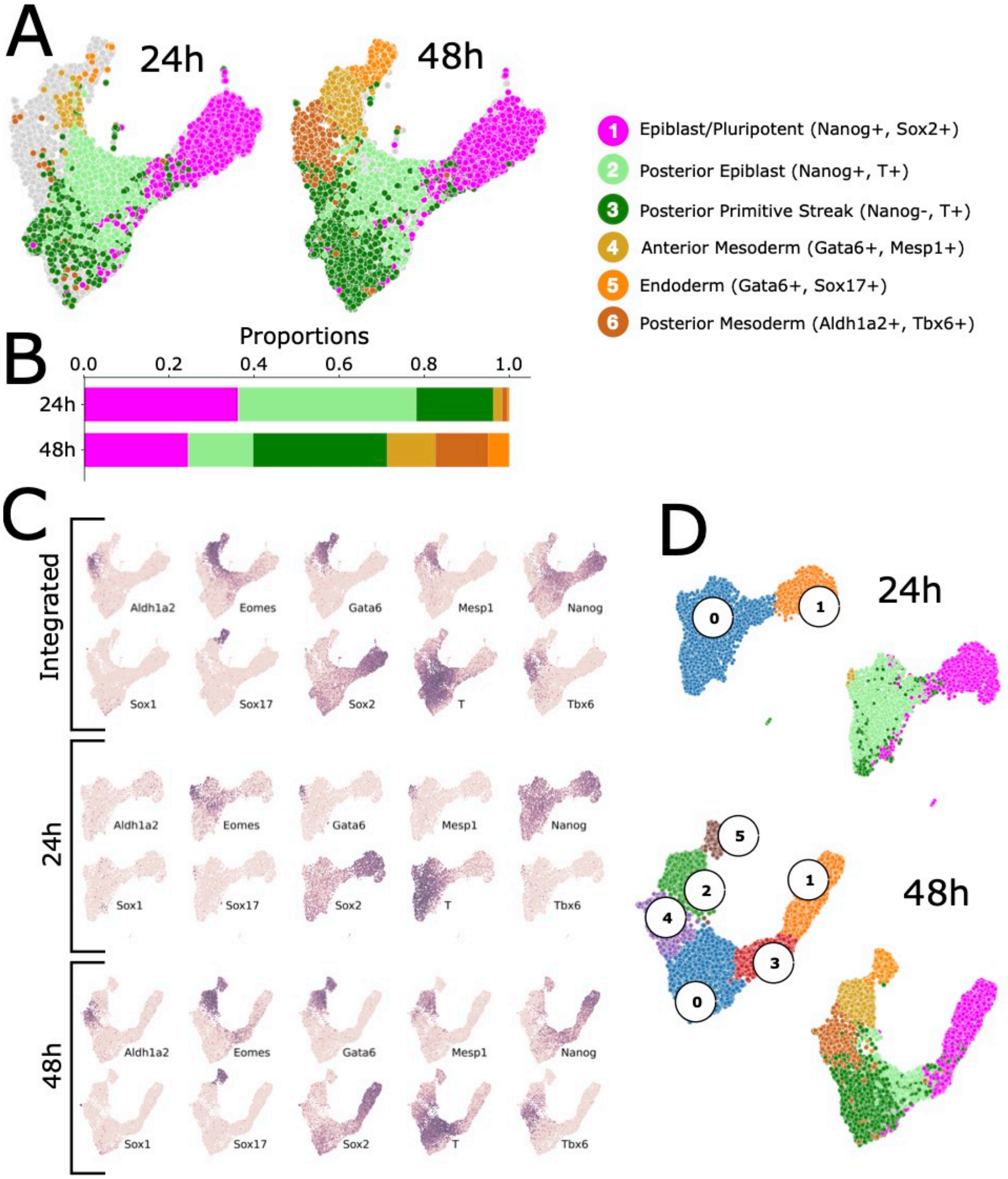


Figure S3: **scRNA-seq analysis of independent gastruloid replicas at 24 hpa and 48 hpa**. A) UMAPs of the integrated analysis of the 24 hpa and 48 hpa datasets with the annotated clusters. B) Proportion of cells per annotation and time point. C) UMAPs of the integrated and independent analysis at 24 hpa and 48 hpa of the dataset, with the marker genes expression color-coded. D) UMAPs of the independent analysis per time-point, showing the leiden clustering and the automatic annotation method.

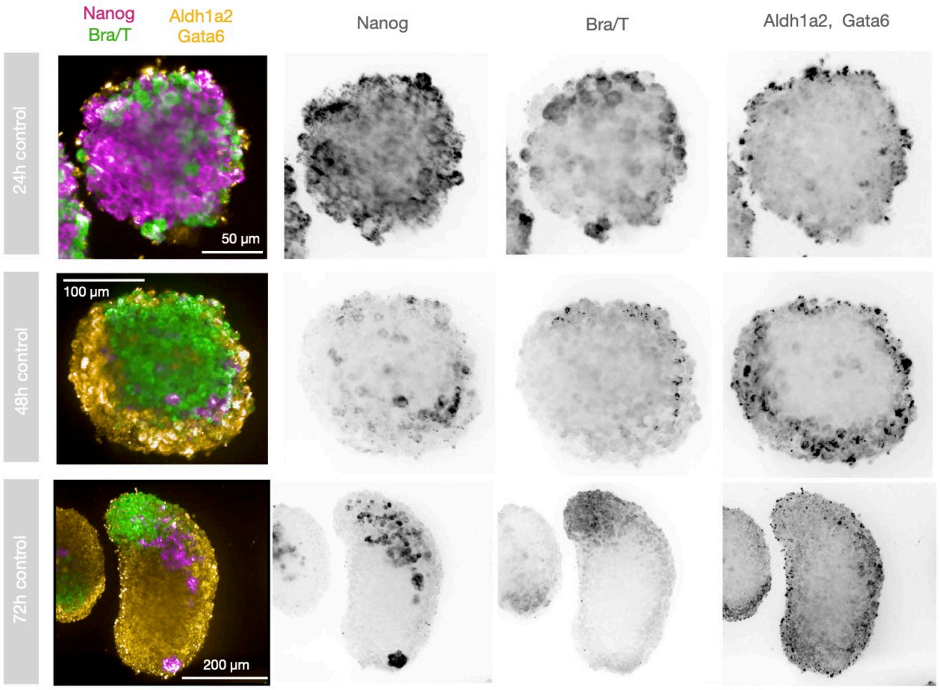


Figure S4: **HCR stainings of gastruloids**. Second replicate as a control. The dynamic range is the same in all images. The images correspond to a maximum intensity projection. T: 488 nm, Nanog: 647 nm, Aldh1a2, Gata6: 546 nm.

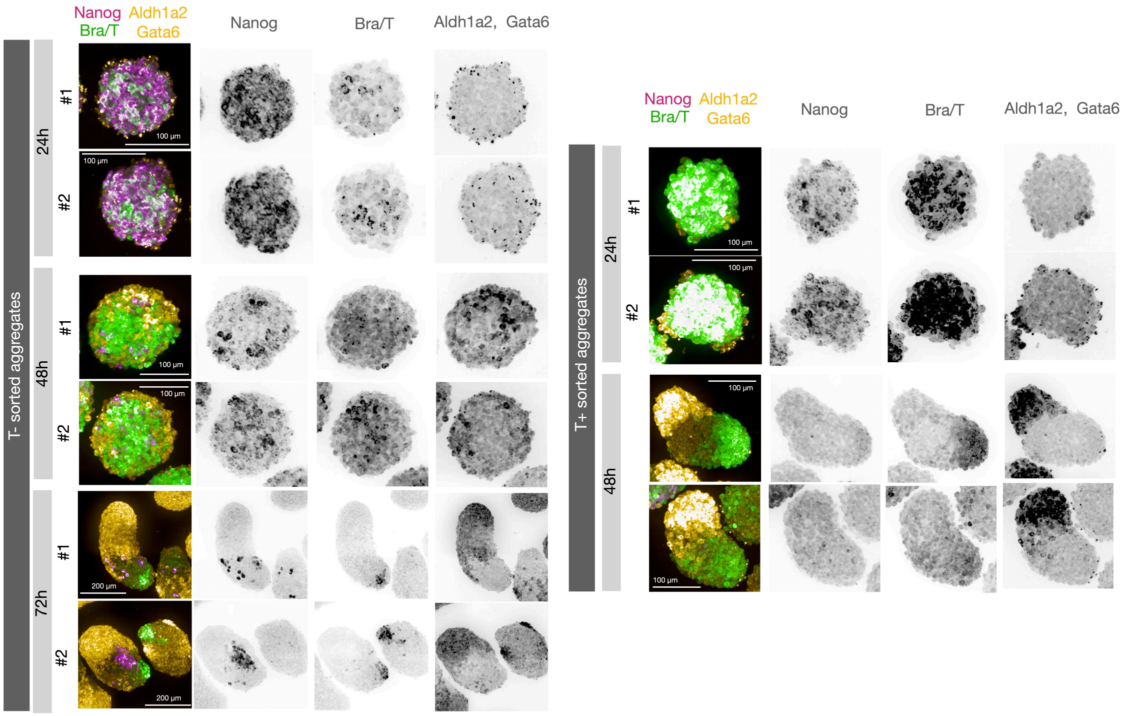


Figure S5: **HCR stainings of T- and T+ aggregates**. The dynamic range is the same in all images and the staining protocol was applied in parallel to the T+ and T- aggregates. The images correspond to a maximum intensity projection. T: 488 nm, Nanog: 647 nm, Aldh1a2, Gata6: 546 nm. No clear signal of the markers could be observed for 72 hpa T+ aggregates.

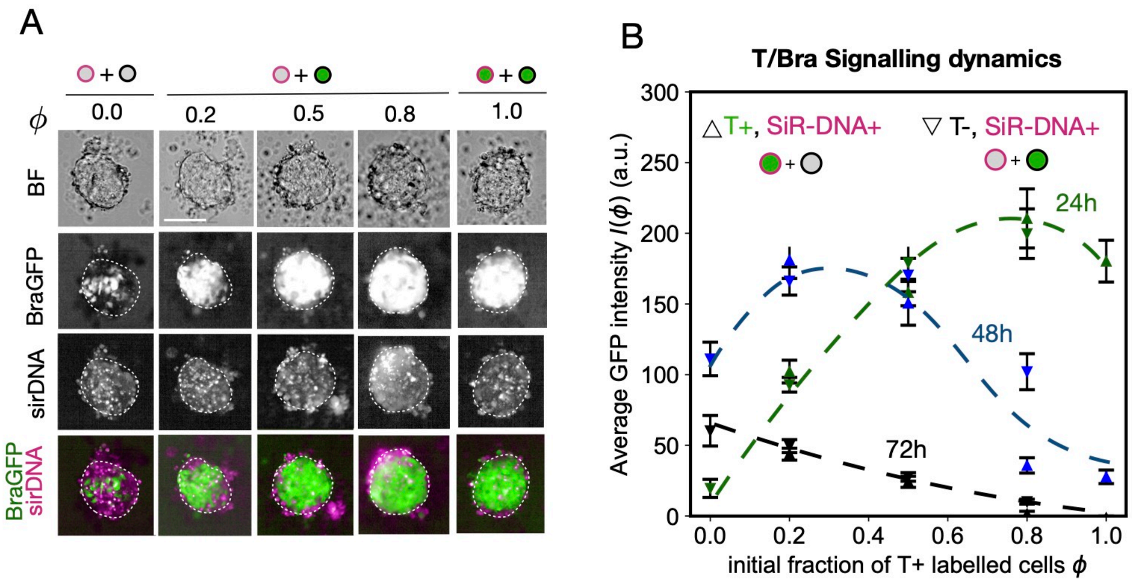


Figure S6: **3D mESC aggregates at 24 hpa for different initial fractions** $\phi$. A) Confocal imaging of 3D mES cell aggregates. The panel shows brightfield (BF), T (GFP) and SiR-DNA signals. The images correspond to maximum intensity projections. Scale bar: 100 $\mu$m. B) Average GFP intensity $I(\phi)$ versus the initial fraction of T+ labelled cells $\phi$ adding SiR-DNA to the T+ (upper triangles) or the T- (lower triangles) populations. T dynamics in the aggregates remained unaffected upon the addition of SiR-DNA. Dashed lines are only a guide to the eye. Error bars correspond to SEM.

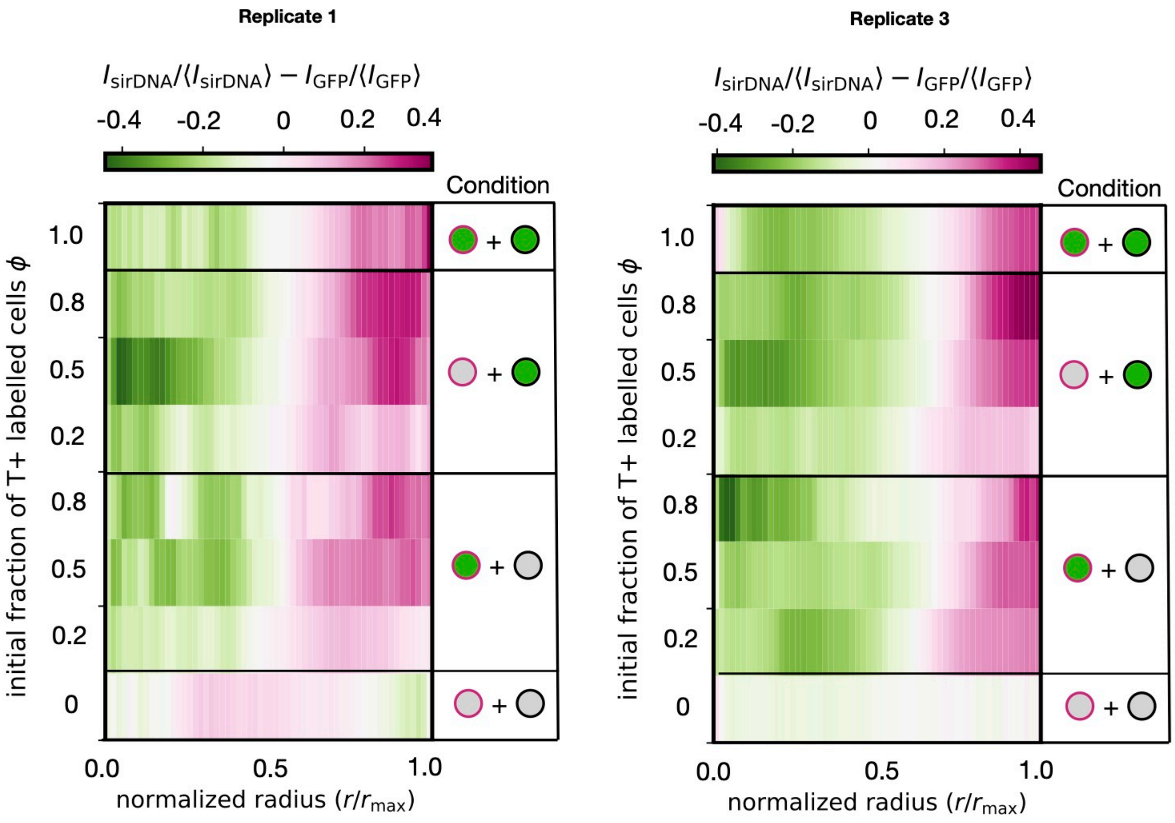


Figure S7: **Radial profiles mESC aggregates**. Colormaps of the radial profile for different starting fractions of T+ ($\phi$) and different conditions. Replicates 1 and 3 are shown, while replicate 2 is shown in Fig. 3D. The replicates were done on different days using different batches of cells.

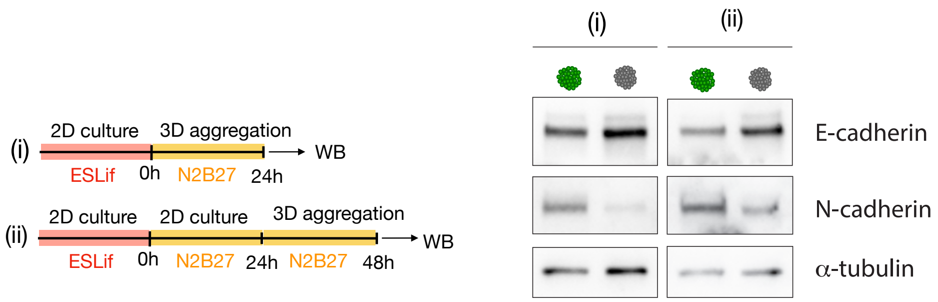


Figure S8: **Cadherin protein levels in T+ and T- aggregates**. FACS sorted T+ and T- aggregates (600 cells) were cultured for 24 hpa (i) or 48 hpa (ii) in N2B27 prior to the lysate. Western blots showed differential protein levels of E and N-cadherin. N- and E-cadherin levels were higher in T+ and T- aggregates, respectively.

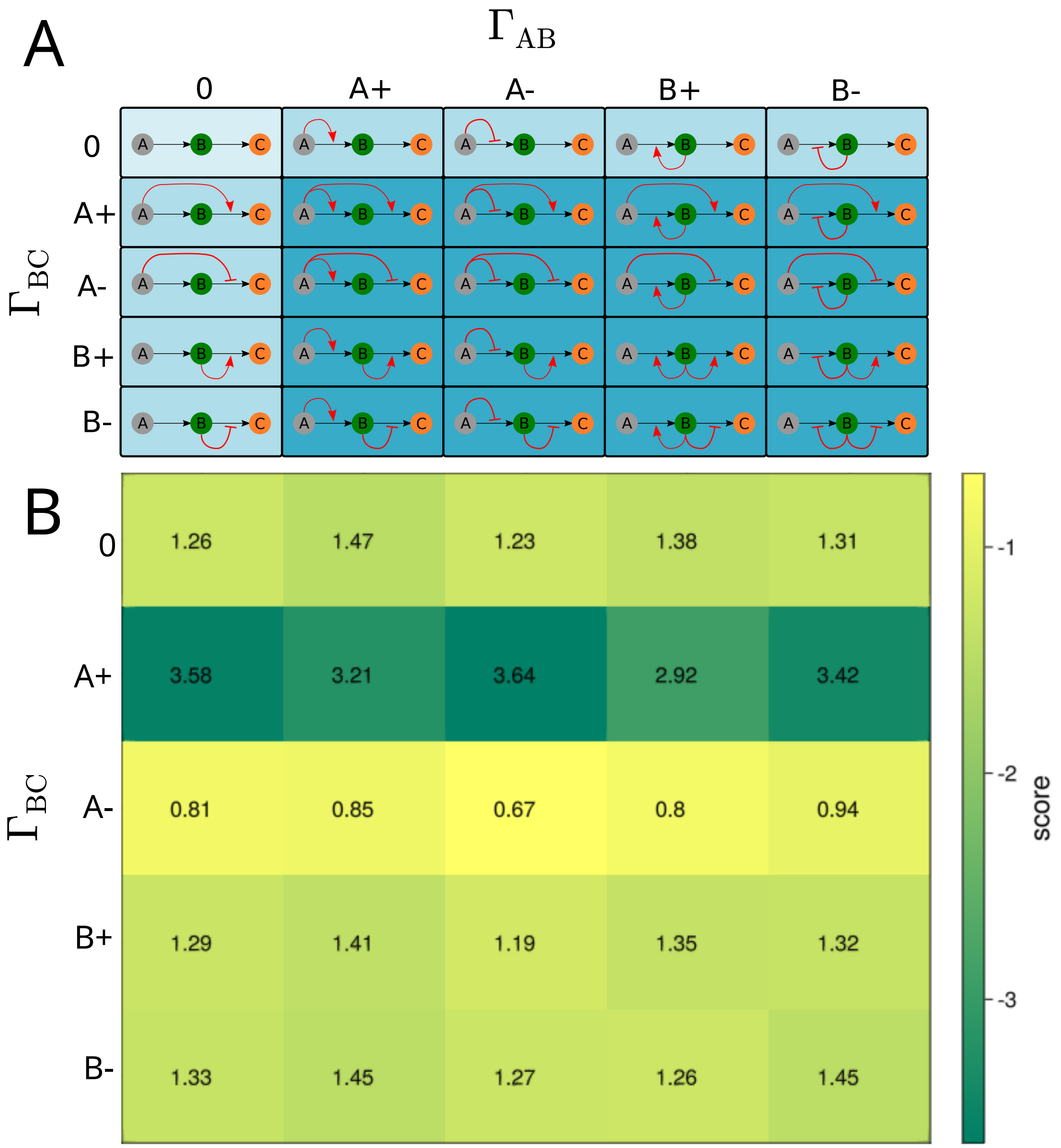


Figure S9: **Cell-cell signalling models**. A) Possible cell-cell signalling models of up to two distinct regulation arrows, on the transition ratios between cell states for the three-state model. B) Loss score of the fit to the data for each of the models described in panel (A). The lower the score, the better the fit to the model. "A+/A-" or "B+/-" indicates that states A or B up-regulate/down-regulate a certain cell fate transition. "0" indicates that a certain cellular state is not regulating a cell fate transition.

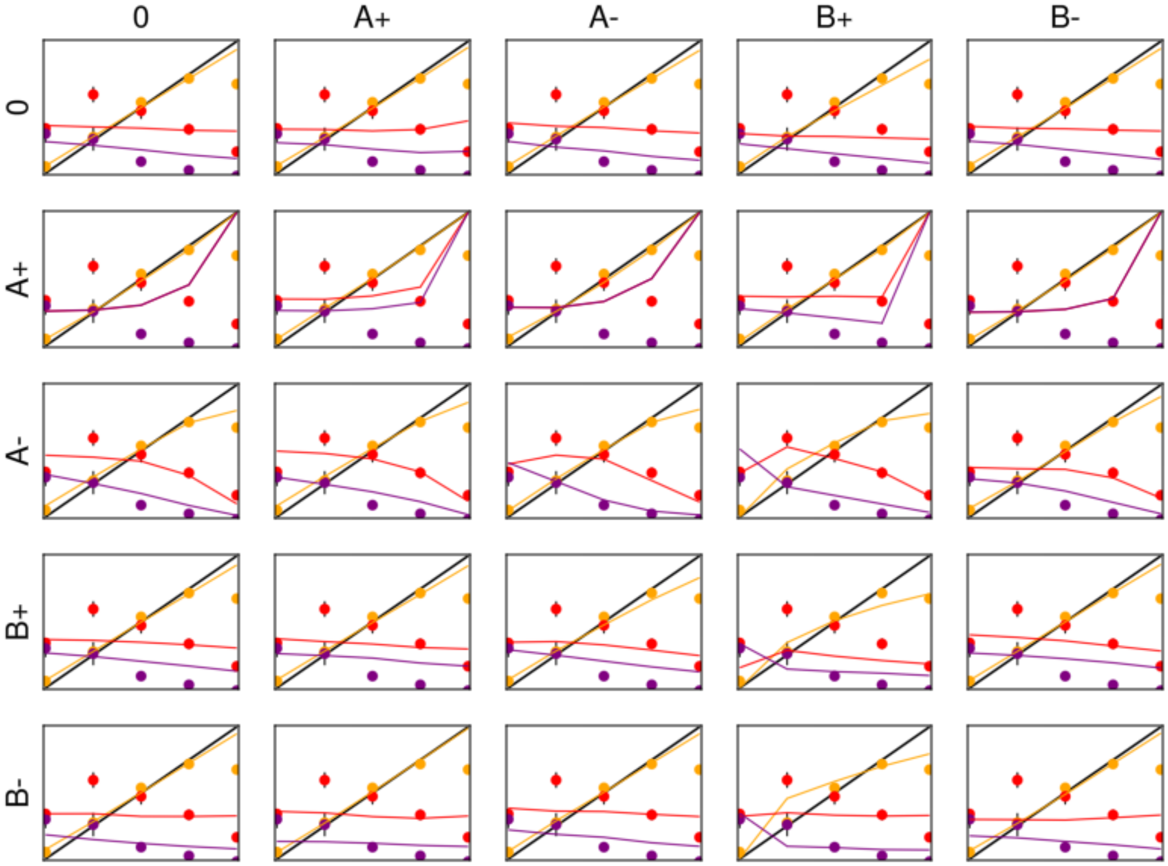


Figure S10: **Best fitting over possible cell-cell communication models**. Solid lines displaying the trends of the best parameter fitting for each model, described in Supplementary Figure S9. Filled circles correspond to the experimental data in Fig. 1D (Yellow: 24 hpa, Red: 48 hpa and Purple: 72 hpa).

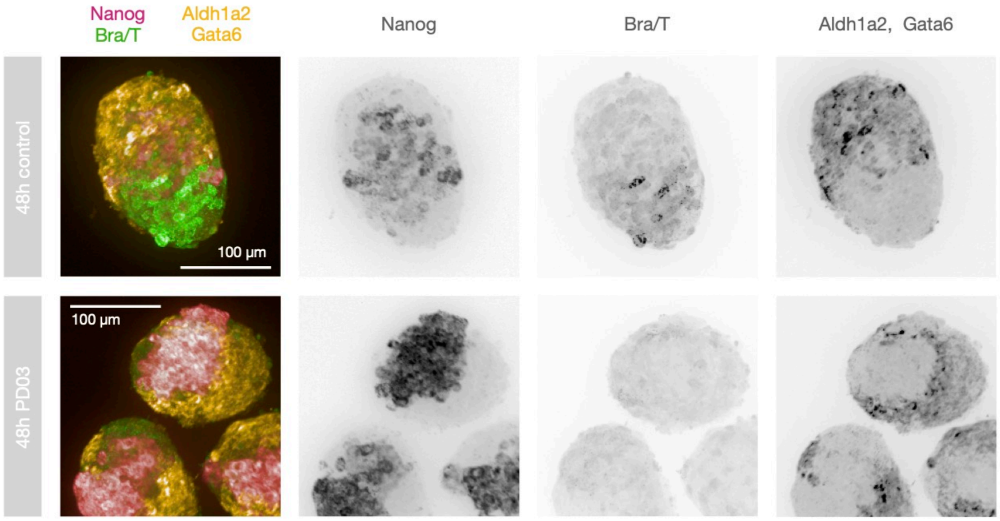


Figure S11: **HCR stainings of 48 hpa cell aggregates for the case $\phi(0) = 0.5$ and $1\mu$M PDO3 inhibition from 24 hpa to 48 hpa**. We observe the lack of Bra/T signal in the inhibited aggregates, showing that the A to B transition is halted upon PDO3 treatment. T: 488 nm, Nanog: 647 nm, Aldh1a2, Gata6: 546 nm.

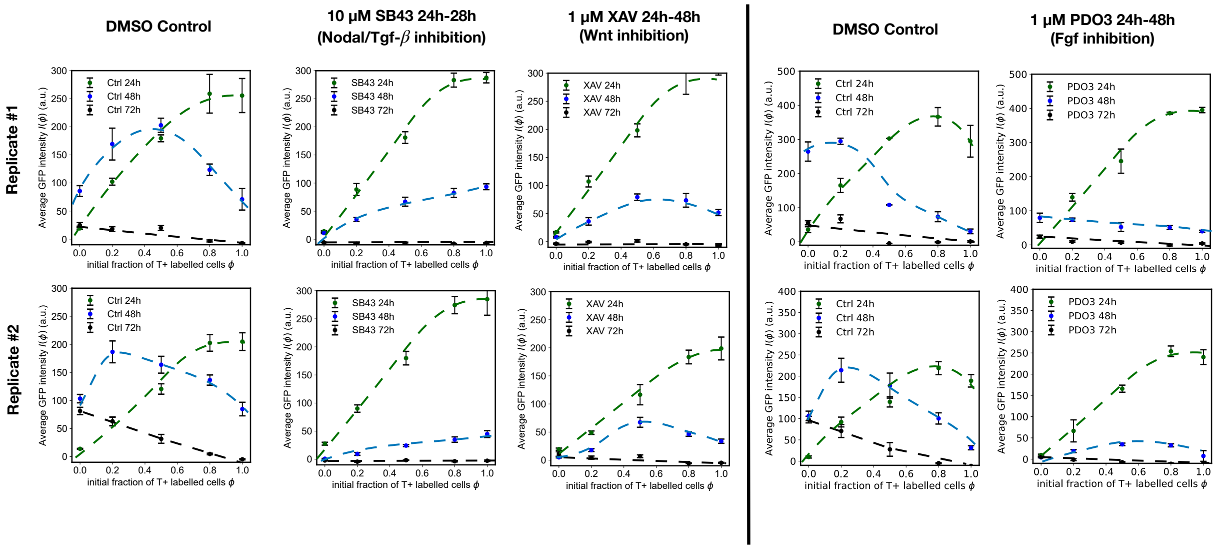


Figure S12: **Effect upon the inhibition of Wnt, Nodal/Tgf-$\beta$ and Fgf pathways on the evolution of the fraction of cells expressing T at 24, 48 and 72 hpa**. The pathways were inhibited using 1 $\mu$M XAV, 10 $\mu$M SB43 and 1 $\mu$M PDO3, respectively. The different inhibitors were incubated from 24 to 48 hpa. Mean $\pm$ SEM. The replicates for Fgf inhibitions were conducted on different days than the Wnt and Nodal/Tgf-$\beta$ inhibitions. Dashed lines are only a guide to the eye.

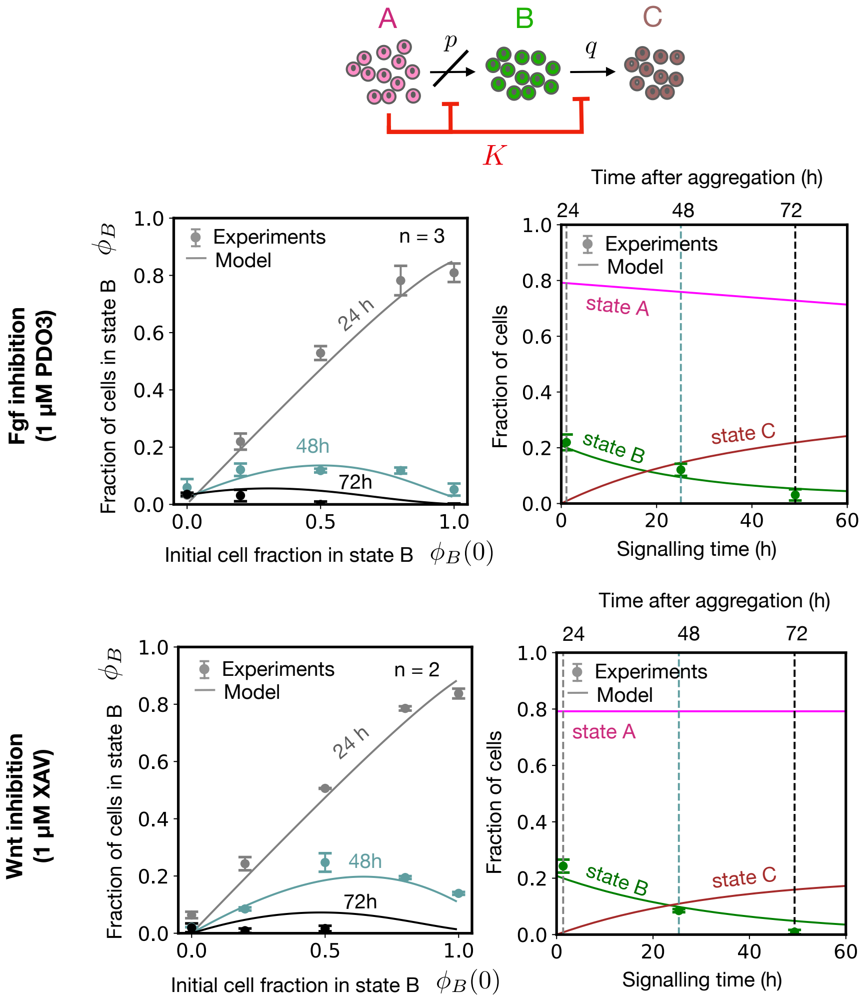


Figure S13: **Effect upon the inhibition of Wnt and Fgf pathways on the evolution of the fraction of cells expressing T at 24, 48 and 72 hpa**. (Left) Cell proportion plots as a function of time. (Right) Fractions of cells $A$, $B$ and $C$ as a function of time. For both cases, solid lines correspond to fits to the model. The Wnt pathway was inhibited with 1 $\mu$M XAV and the Fgf pathway was inhibited with 1 $\mu$M PDO3. Both inhibitors were incubated from 24 to 48 hpa. Error bars correspond to 2 $\times$ SEM over different replicates. For the case of Wnt inhibition, the fit with $p = 0$ is shown; however, for the case of Fgf inhibition, the best fit for small non-zero $p$ is shown (see Table S3)

| Annotation | Genes |
|---|---|
| Epiblast/Pluripotency | Nanog+, Sox2+ |
| Posterior Epiblast | Nanog+, T+ |
| Posterior Primitive Streak | Nanog-, T+ |
| Anterior Mesoderm | Eomes+, Gata6+, Mesp1+ |
| Posterior Mesoderm | Aldh1a2+, Tbx6+ |
| Endoderm | Eomes+, Gata6+, Sox17+ |

Table S1: **Annotation markers**.

| Antibody | Company | Catalogue number | Dilution |
|---|---|---|---|
| anti-E-Cadherin | Cell Signaling | 3195 | 1:2000 |
| anti-N-cadherin | Abcam | ab18203 | 1:2000 |
| anti-$\alpha$-catenin | Merck (Sigma) | T6199 | 1:1000 |

Table S2: **Primary antibodies used for the Western Blots**.

| Condition | $n$ | $p\ (h^{-1})$ | $q\ (h^{-1})$ | $K$ | $T_{24}\ (h)$ |
|---|---|---|---|---|---|
| Control | 4 | $0.14 \pm 0.01$ | $0.071 \pm 0.003$ | $0.17 \pm 0.02$ | $5.1 \pm 0.5$ |
| PDO3 | 4 | $0.006 \pm 0.001$ | $0.15 \pm 0.01$ | $0.30 \pm 0.04$ | $1.09 \pm 0.06$ |
| | 3 | 0 (fixed) | $0.4 \pm 0.1$ | $0.10 \pm 0.07$ | $0.6 \pm 0.1$ |
| XAV | 4 | $(8.8 \pm 0.2) \cdot 10^{-5}$ | $0.088 \pm 0.002$ | $0.40 \pm 0.03$ | $1.36 \pm 0.03$ |
| | 3 | 0 (fixed) | $0.088 \pm 0.002$ | $0.40 \pm 0.03$ | $1.4 \pm 0.1$ |
| SB43 | 4 | $(7.2 \pm 0.1) \cdot 10^{-5}$ | $0.072 \pm 0.001$ | $1.7 \pm 0.2$ | $1.6 \pm 0.1$ |
| | 2 | 0 (fixed) | $0.0660 \pm 0.0004$ | $\infty$ (fixed) | $1.5 \pm 0.1$ |

Table S3: **Fitted model parameters in the different conditions.** The value $n$ indicates the number of fitted parameters.

Table S4: Main agent-based model parameters. $^\star$: Orientative cell-division timescale estimated from Ref. (*36*). $^\dagger$: Experimentally obtained parameters in this work.

| Parameter | Symbol | Value |
|---|---|---|
| Time step | $\tilde{\Delta t}$ | 0.02 |
| Characteristic length | $\tilde{r}$ | 1 |
| Average cell division time for state $A$ | $\tilde{\tau}_{\text{div},A}$ | 350$^\star$ |
| Average cell division time for state $B$ | $\tilde{\tau}_{\text{div},B}$ | 350$^\star$ |
| Average cell division time for state $C$ | $\tilde{\tau}_{\text{div},C}$ | 750 |
| SD of cell division time | $\tilde{\sigma}_{\text{div}}$ | 0.5 |
| Effective friction coefficient | $\tilde{\lambda}$ | 1 |
| Cell-ECM/cell-cell friction parameter | $\alpha$ | 0.9 |
| $A \rightarrow B$ transition rate | $\tilde{p}$ | 0.009333$^\dagger$ |
| $B \rightarrow C$ transition rate | $\tilde{q}$ | 0.004667$^\dagger$ |
| Cell division relaxation time | $N_{rel}$ | 150 |
| Feedback parameter | $K$ | 0.2$^\dagger$ |

Table S5: Homotypic and heterotypic interaction parameters. Matrices are symmetric; only the upper triangle is shown.

Attraction force $\tilde{F}_{\text{att}}^{(i,j)}$

| | A | B | C |
|---|---|---|---|
| A | 3.6 | 4.0 | 2.4 |
| B | – | 6.4 | 1.6 |
| C | – | – | 3.0 |

Repulsion force $\tilde{F}_{\text{rep}}^{(i,j)}$

| | A | B | C |
|---|---|---|---|
| A | 9.0 | 3.0 | 3.0 |
| B | – | 6.0 | 3.0 |
| C | – | – | 6.0 |

Force range $\mu_{ij}$

| | A | B | C |
|---|---|---|---|
| A | 2.0 | 1.82 | 2.0 |
| B | – | 2.0 | 2.0 |
| C | – | – | 2.14 |
